# Semi-Automated High Content Analysis of Pollen Performance Using TubeTracker

**DOI:** 10.1101/2024.11.21.624782

**Authors:** Sorel V. Yimga Ouonkap, Yahir Oseguera, Bryce Okihiro, Mark A. Johnson

## Abstract

Pollen function is critical for successful plant reproduction and crop productivity and it is important to develop accessible methods to quantitatively analyze pollen performance to enhance reproductive resilience. Here we introduce TubeTracker as a method to quantify key parameters of pollen performance such as, time to pollen grain germination, pollen tube tip velocity and pollen tube survival. TubeTracker integrates manual and automatic image processing routines and the graphical user interface allows the user to interact with the software to make manual corrections of automated steps. TubeTracker does not depend on training data sets required to implement machine learning approaches and thus can be immediately implemented using readily available imaging systems. Furthermore, TubeTracker is an excellent tool to produce the pollen performance data sets necessary to take advantage of emerging AI-based methods to fully automate analysis. We tested TubeTracker and found it to be accurate in measuring pollen tube germination and pollen tube tip elongation across multiple cultivars of tomato.

Graphical Abstract
Graphical user interface of TubeTracker showing all supported functionalities.

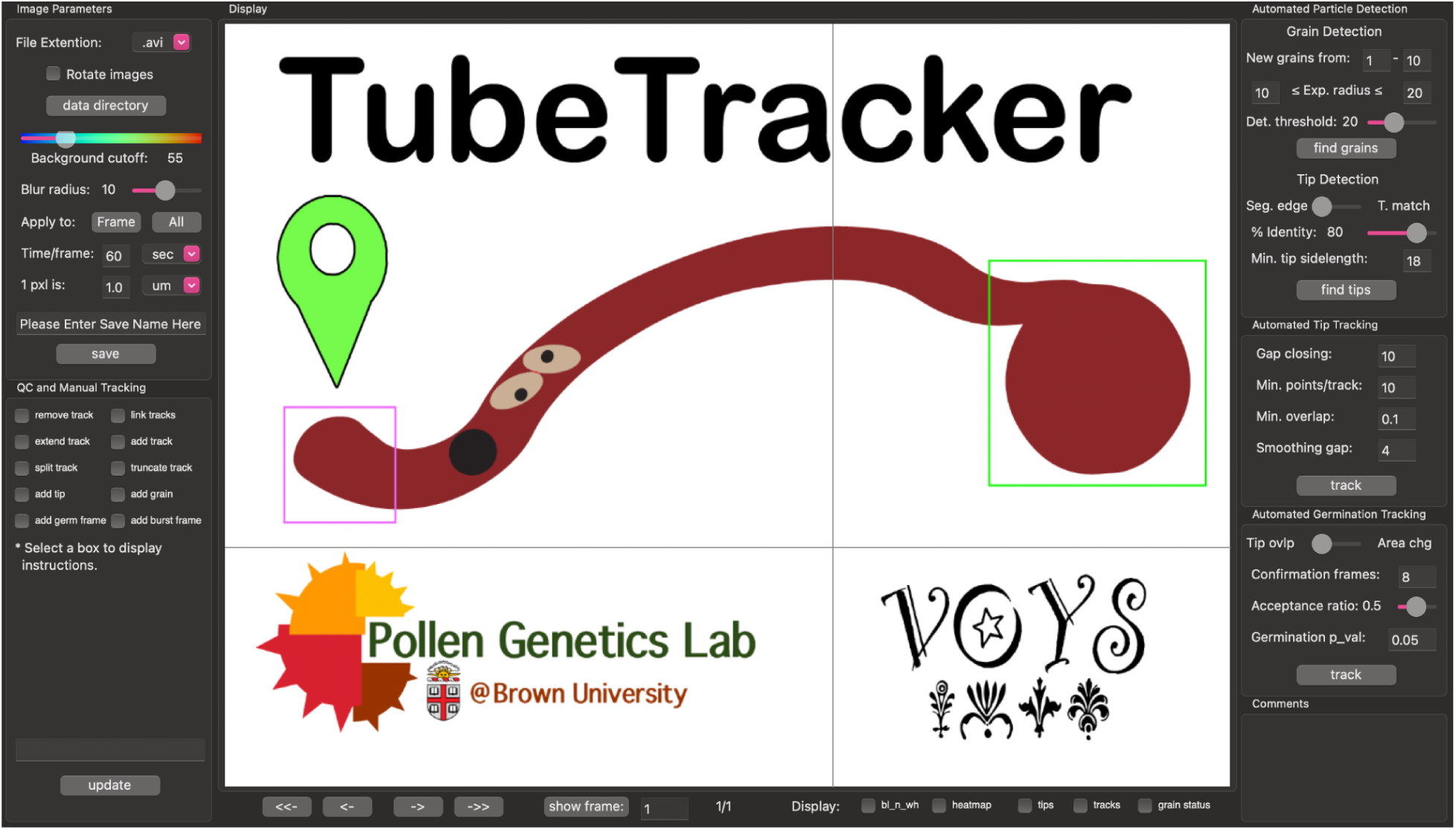

## Introduction

### Research into pollen grain and tube function is important for food sustainability

Plant sexual reproduction is important for the production of seeds, kernels, and fruits, which together are the basis of agriculture and are essential to feed an ever-growing world population. This key stage of the flowering plant life cycle depends on a pollen grain, the male gametophyte, which germinates a pollen tube that burrows through pistil tissue via tip extension until it reaches an ovule in the ovary. After recognition by synergid cells of the female gametophyte, the pollen tube undergoes cell rupture to deliver two sperm cells for double fertilization of the egg and central cell, leading to seed development [1]. Because each fertilized ovule develops into one seed, the number of seeds is directly proportional to pollen performance, which is defined as the ability of pollen grains to germinate a pollen tube that successfully extends to and ruptures in an ovule. Pollen performance can also influence fruit biomass, as we have shown that for cultivated tomato (*Solanum lycopersicum*, hereafter referred to as tomato), the number of seeds produced in a fruit is directly proportional to the fruit’s biomass [2]. Furthermore, some cultivars that maintain fruit and seed production at high temperature also maintain high pollen tube performance in the pistil [2]. This indicates that studying the pollen tube growth phase is critical to improving crop productivity under the influence of different external and internal stimuli.

Unfortunately, the pollen tube growth phase is highly vulnerable to multiple environmental stresses including drought [3–12] and rising temperatures ([2,13–21]). As a result, considerable efforts have been made to understand and improve reproductive output under extreme conditions. One promising approach is to study natural variation present in crops that are resistant to extreme conditions in order to incorporate these causal variants into susceptible but commercially important crops. This will require phenotypic analysis of plants to identify the defective phases of reproduction, identification of molecular mechanisms that confer resistance, and finally targeted genomic improvements that enhance yield under stress while also maintaining other desired traits.

### Phenotypic analysis of pollen germination and elongation is currently limited

As a key contributor to reproductive success, pollen development/production has been the subject of extensive research in part because it is relatively easy to determine the number of pollen grains that have been produced. On the other hand, the absence of proper methods to dissect each stage of pollen tube growth through accurate quantitative phenotypic analyses has been an obstacle to progress in understanding how pollen performance is affected by biotic and abiotic factors such as heat stress.

In vitro pollen performance is defined as the ability of pollen tubes to germinate and rapidly elongate a pollen tube while avoiding pollen tube burst. Pollen performance can be divided into its main components: 1) pollen grain germination, 2) pollen tube tip elongation rates, and 3) pollen grain and tube viability (hereafter referred to as pollen survival rate, Sup. Fig S1). Currently, pollen performance is only measured using end-point analyses which quantify the fraction of pollen that germinates, the length of pollen tubes, and the fraction of pollen that survives after incubation for a set amount of time. These analyses are usually slow and limit the number of samples that can be collected and quantified. Additionally, pollen tube growth is a dynamic process that happens over hours, while end-point analyses only provide a snapshot of the process. To properly quantify pollen performance, we should include time. If time is made a dynamic variable in the analysis of pollen tube growth in vitro, we can determine the specific time of germination, whether germination is synchronous and differs among genotypes and species. We can further determine the extension rate of the tip at various time points and how pollen tube survival rates change over time. The ability to measure these pollen performance parameters using methods with relatively high throughput will open up many possibilities for discovery about genetic variants as well as biotic and abiotic conditions that affect each parameter over time. Finally, time-lapse analysis of the pollen tube opens the door to new statistical analysis such as survival analysis [22–25], which has been a staple approach in other fields [26,27].

Current methods for pollen grain and tube phenotypic analysis are limited. Many approaches have been used to characterize pollen germination [28]. These include manual measurement, often made using Fiji/ImageJ [29,30], as well as Kymograph-based technologies [31–34]. These methods, however, limit the number of pollen tubes that can be analyzed and thus the number of experiments that can be performed on a reasonable timeline. Furthermore, while these manual methods can measure terminal germination percentage, pollen tube length, and survival (the fraction of grains and/or tubes that do not burst/rupture over time), they will miss any changes that happen over time, and they cannot measure important performance parameters like time to germination, extension rates, per-pollen elongation time, and time of pollen tube rupture. These latter parameters can only be obtained via analysis of time-lapse series and may require facilitation of data quantification using semi-automation.

Other approaches have introduced automation to quantify pollen tube performance over time but are limited by either the range of analyses performed, the state of pollen tubes after analysis, or the ease of implementation or transferability of the method. These recent tools include those focusing solely on pollen germination such as ‘pollen_tubes’ (https://github.com/JIC-Image-Analysis/pollen_tubes), which implements an algorithm that detects pollen grains and another that counts all particles in an image. Percent germination is then calculated using the ratio of all particles to pollen grains. This method, however, requires the use of two different microscopes for resolution and does not provide any information about pollen tube elongation. Methods such as ASIST focus on pollen elongation by using iterative subtraction of successive temporal frames to identify the location of pollen tube tips and then connect these to create pollen tube trajectories [28]. This method, however, lacks analysis of pollen germination and the tracking method breaks down if the pollen tube experiences a significant lateral shift in successive frames.

Machine learning combined with object detection offers great potential to automate analysis of pollen performance parameters and tools are being developed to take advantage of this technology (“pollen_cv”, https://github.com/cedarwarman/pollen_cv). While machine learning methods are highly accurate, they suffer from poor transferability between research groups. To implement these tools for analysis of images collected from different imaging platforms, each new user must train the algorithm with an annotated training data set produced using their imaging platform. This process is time-consuming, resource-intensive and will be prohibitive for many potential users/applications. Despite their low transferability, machine learning tools will considerably improve quantification of pollen performance and will be particularly useful for analysis of large numbers of genotypes or for screening of large chemical libraries.

Here, we took advantage of progress made to open-source image analysis modules such as OpenCV [35] and Wxpython [36,37] to create TubeTracker, a comprehensive Python-based software that does not require training of the analysis algorithm, and can be easily implemented and used by any research group to analyze pollen germination, pollen tube elongation, and pollen grain/tube survival. The tool is user friendly, features are clearly defined in our user manual (Sup. File S1) and the data produced can be used to annotate key pollen classes needed to train machine learning based detection algorithms.

## Results

### Brightfield image normalization using horizontal and vertical line detection

To make TubeTracker easily usable across multiple laboratory and imaging platforms, we wanted to normalize all input time-lapse images to generate uniform black and white images containing recognizable pollen grain/tube particles optimized for downstream processing. Our goal was to remove all background pixels and retain only pixels corresponding to pollen grains and tubes. To differentiate pollen pixels from background pixels, TubeTracker implements a segmentation method based on the Sobel operator, which detects particle edges based on pixel contrast variations [38,39]. The Sobel operator allows us to detect particle edges in the horizontal and vertical axes separately. These detected particle edges can then be merged together to reconstruct a high fidelity black and white image in which pixel intensities depend on contrast present in the input image (Fig 1A, 1B; segmented image is color-coded according to pixel intensity).

**Figure 1:**
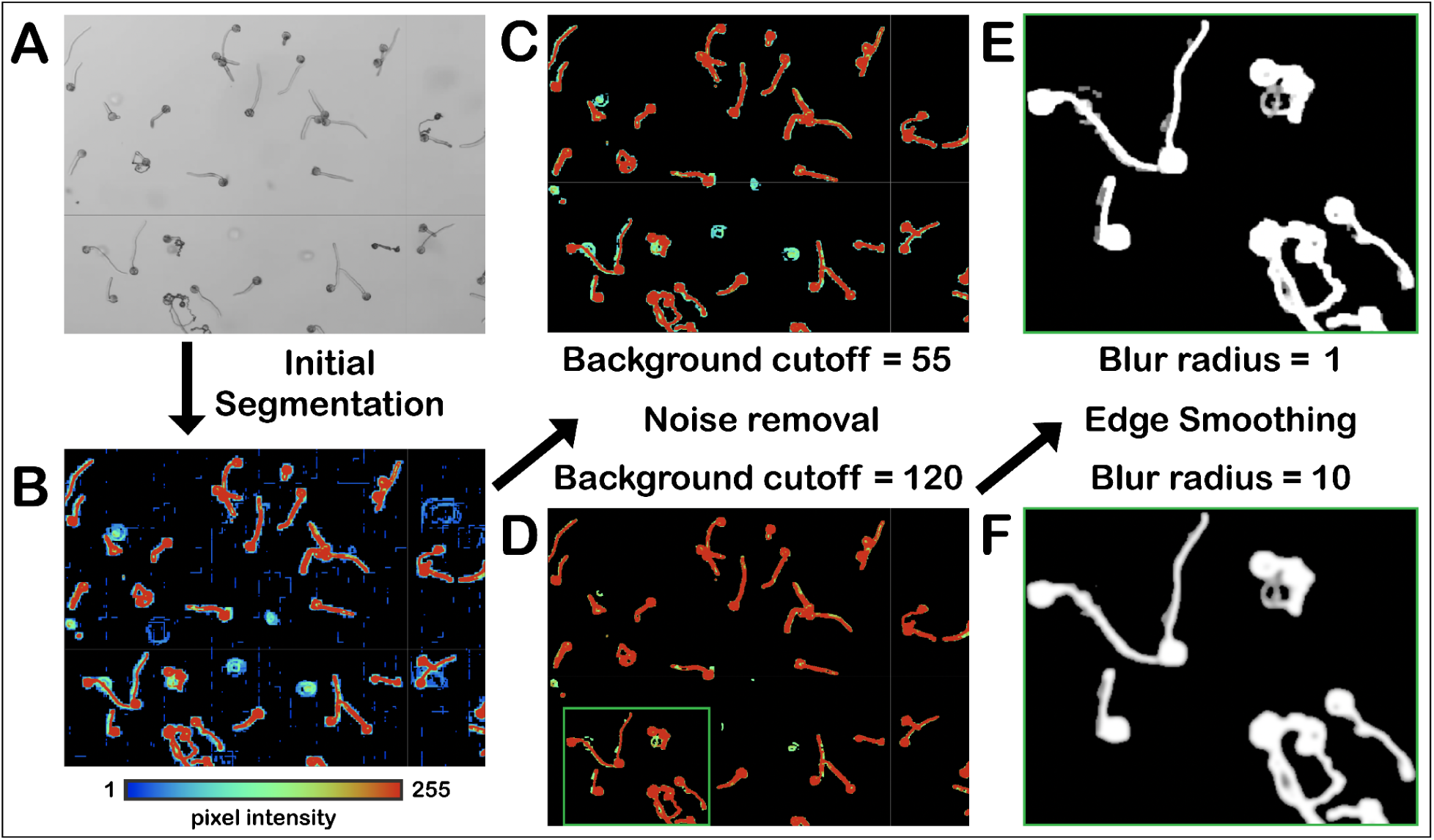
TubeTracker’s uses segmentation to create binary images from each frame. (**A)** Representative input frame collected using brightfield. (**B)** Initial segmentation resulting from horizontal and vertical line detection followed by pixel coloring according to pixel intensity. (**C, D)** Particle segmentation can be tuned using different background intensities. (**E, F)** Effects of different blur radii on particle segmentation. Note that only the green area from D is shown to highlight the effects of this filter.

Due to the high sensitivity of the Sobel operator to contrast changes, a significant amount of low intensity noise is also detected (Fig 1B). To improve the quality of the segmented image, TubeTracker offers two filters. The first filter removes all pixels with intensity values below a user-defined threshold (Fig. 1C, 1D, “Background cutoff”, Sup. File S1, pp 13-14). The application of this background filter produces particles with grainy edges that are suboptimal for downstream processing (Fig 1E). To improve downstream processing, TubeTracker offers a second filter that implements opencv’s gaussian blurring method for which the user must provide a blur radius of action (“Blur radius”, Sup. File S1, pp 13, 15). This final processing step produces time-lapse frames that contain particles with smoother and uniform edges and that are optimal for downstream analysis steps (Fig. 1E vs. 1F; Note that a blur radius of 1 indicates no blurring, Fig. 1E). These user-defined processing steps can be tailored to each frame or to a time-lapse series.

### Particle Detection

To track pollen grain germination, pollen tube elongation and pollen grain/tube survival over time, five different types of particles must be detected from a set of time-lapse images, 1) ungerminated pollen grains, 2) germinated pollen grains, 3) burst pollen grains, 4) pollen tube tips, and 5) burst pollen tube tips (annotated using red, green, blue, pink, and cyan boxes respectively, Sup. Fig. S1). TubeTracker automatically detects ungerminated pollen grains, germinated pollen grains, and pollen tube tips (Fig. 2A, Fig. 3A) and facilitates manual detection of all five particle types (Fig. 2B).

**Figure 2:**
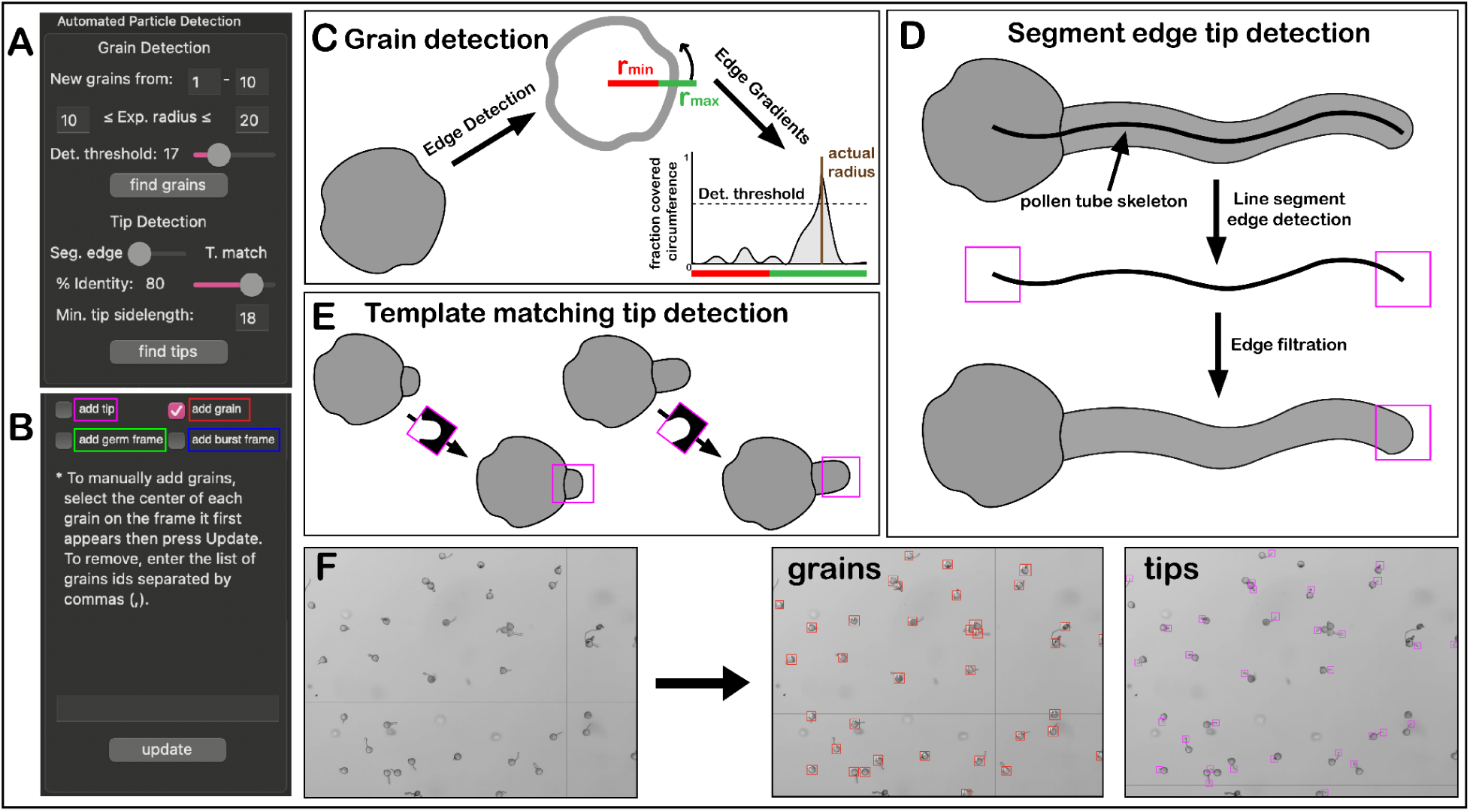
Detection of pollen grains and pollen tube tips using TubeTracker: (**A)** The graphical user interface allows the user to adjust parameters to detect pollen grains and pollen tube tips. (**B)** The graphical user interface showing the user-controlled features that manually complement detection of important particles. (**C)** Illustration of the algorithm used to detect grains. (**D)** Illustration of the segment edge tip detection algorithm used to detect pollen tube tips. **(E)** Illustration of the template matching tip detection algorithm used to detect pollen tube tips. (**F)** Representative images showing the results of automatic grain and tips detection.

**Figure 3:**
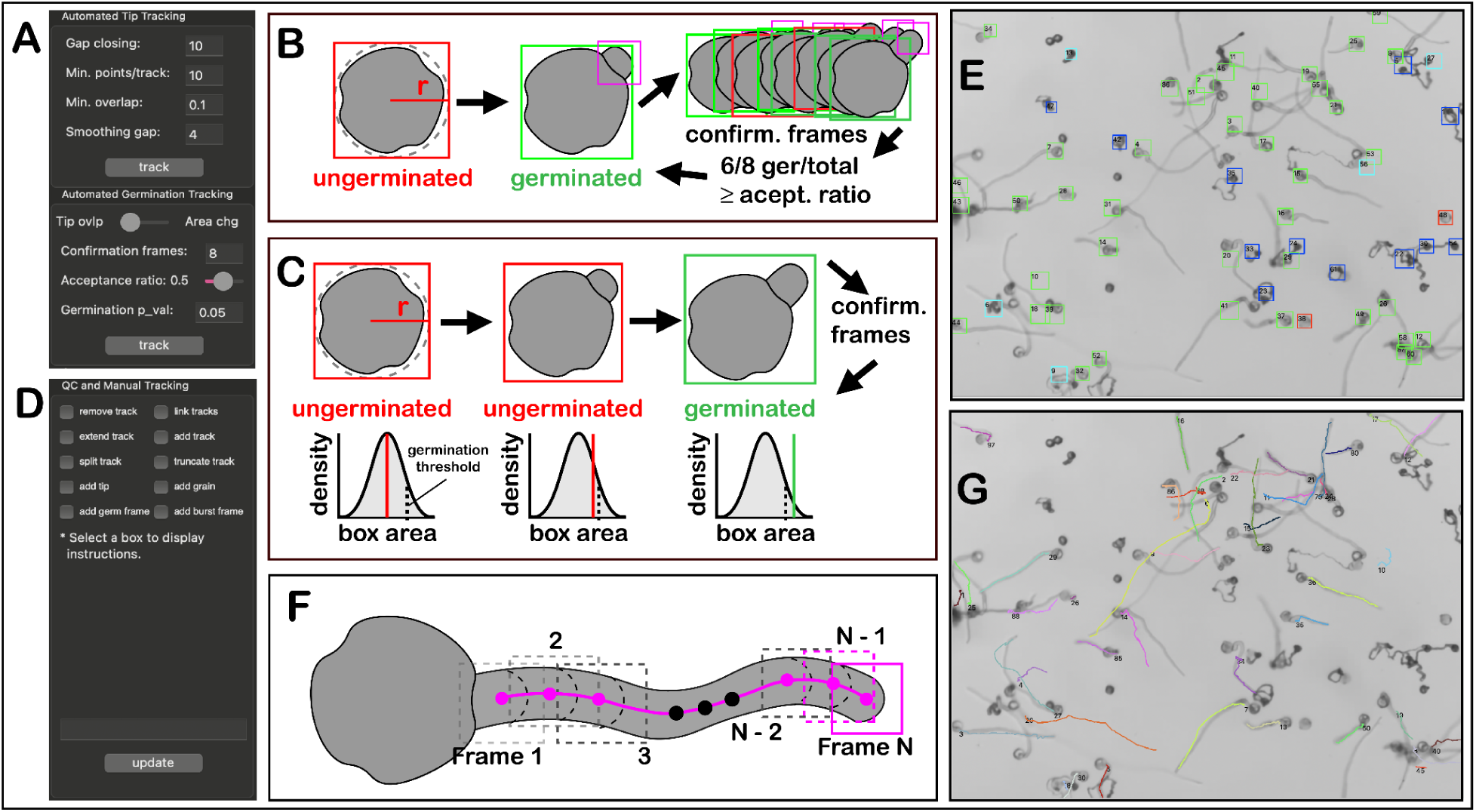
TubeTracker’s quantification of germination, survival and tip elongation: (**A)** The graphical user interface showing the user-controlled parameters used for automated germination tracking and automated elongation tracking. (**B)** Illustration of the tip-overlap algorithm used to detect germinated pollen grains. (**C)** Illustration of the secondary algorithm used to detect germination pollen grains based on area change. (**D)** The graphical user interface showing the features that can be used to manually track germination, survival and elongation of tracks and to complement automated tracking. (**E)** Visual results of semi-automated tracking of pollen germination and survival. Red: ungerminated pollen grains, Blue: burst pollen grains, Green: germinated and actively elongating pollen tubes, Cyan: germinated and burst pollen tubes. (**F)** Illustration of the algorithm used to track the elongation rate of pollen tube tips. (**G)** Visual results of semi-automated tracking of pollen tube elongation. Each pollen tube is assigned a random color.

To detect pollen grains (which are assumed to be ungerminated at first detection), three variables must be provided by the user (see provided manual for instructions, Sup. File S1) – the range of frames to be used to detect pollen grains (“New grains from”), the expected range of grain radius (“Exp. radius”, in pixels), and a grain detection threshold “Det. threshold” (Fig. 2A, top). To detect pollen grains using these variables, TubeTracker relies on the roundness of pollen grains and their edge gradients. First, the edges of each particle in black and white frames are extracted. Edge gradients are then calculated using the fraction of the circumference covered by detected pixels along a hypothetical circle. If the fraction covered is above a user-set threshold (“Det. threshold”, Fig. 2A), and the gradient max is within the grain radius range (“Exp. radius”, Fig. 2A), then the particle is assumed to be a pollen grain (Fig. 2C). This algorithm is implemented using OpenCV’s Hough circle detection algorithm [35] and allows for detection of most grains present throughout the time-lapse series (Fig. 2F). Mistakes resulting from automated grain detection can be fixed using manual addition of grains that were missed and removal of noise that was defined as grains following provided instructions (“add grain”, Fig. 2B, Sup. File S1, pp. 19-20).

To automatically detect pollen tube tips, TubeTracker can implement one of three user-selected methods. The default and recommended method, called “segment edge tip detection”, is accurate, has a fast processing time, and is preferred when input images have low background and low noise. This method reduces pollen particles to line segments (called skeletons), identifies the endpoints of each line segment and then filters these endpoints to differentiate the tip-end from the grain-end of the skeleton (Fig. 2D). In cases where input images have high noise or background, it is recommended that the user choose the “tip template matching” method to define pollen tube tips (Fig. 2E). In this case, TubeTracker identifies instances of pre-defined tip templates in a given image and filters them according to their percent resemblance to the template (Fig. 2E). This requires two user provided variables (“Min. tip sidelength” and “% identity”, Fig. 2A). Finally, in addition to grain and tips that are detected automatically (Fig. 2), the user can also manually add or remove them using the “add tip” function (Fig. 2B). Other pollen particles important for analysis of pollen survival (such as burst pollen grains and burst pollen tubes) can be manually detected using “add burst frame” (Fig. 2B).

### Automated tracking of pollen germination and elongation

Following pollen grain and pollen tube tip particle detection, TubeTracker can automatically extract key fitness features such as pollen germination and elongation over time using a few user-defined parameters (Fig. 3A). To automatically track germinated grains, TubeTracker implements one of three user-selected methods. The first method termed “grain to tip overlap” searches for overlaps between a detected/ungerminated pollen grain and a tip (Fig. 3B). Once this overlap is established, and before germination is confirmed, TubeTracker checks over a set number of user-defined future frames (“Confirmation frames”, Fig. 3A) to validate the germination event by ensuring that the ratio of the number of frames in which the grain remains germinated over the total number of frames used is at or above another user-defined threshold (“Acceptance ratio”, default = 50%, Fig. 3A, 3B). The “grain to tip overlap” method is the default algorithm, is faster, and is recommended in cases where automatic/manual tip detection is complete across all/most frames.

The second method for defining germination that can be implemented by TubeTracker (“bounding box area change”) tracks changes of the area of the grain bounding box against a null distribution built from the size of all detected grains (Fig. 3C). Once this bounding box area passes a user-defined threshold (“Germination p_val”, Fig. 3A) and persists above the “Acceptance ratio” when the user-defined “Confirmation frames” are analyzed, the grain is considered germinated (Fig. 3C). This method is the preferred method when tip detection is inconsistent or absent.

The user has the option to manually define pollen tube germination selecting the “add germ frame” option (Fig. 3D, see provided manual for instructions, Sup. File S1). The user can also use this feature to refine automated detection of germination. The final result is a time-lapse series of images each containing the status of each pollen coded in the color of its bounding box (red: ungerminated, blue: dead grain, green: germinated, cyan: dead pollen tube, Fig. 3E, Sup. Fig. S1).

To track pollen tube tip elongation (length over time), TubeTracker implements a method based on the multi-object tracker package motpy [40,41]. To track tips over time, their location on each frame – previously identified automatically and/or manually (Fig. 2B, 2D, 2E) – are linked to each other based on a user-defined intersection over union area ratio (i.o.u ratio,“Min. overlap”, Fig. 3A, 3F). This automated tracking, which can be refined manually by the user (Fig. 3D, see provided manual for instructions, Sup. File S1 pp. 24-26), produces frames in which the current and/or past positions of pollen tube tips are highlighted (Fig. 3G).

### TubeTracker provides accurate measurements of pollen grain germination and pollen tube elongation

To determine if automated measurements made using TubeTracker are accurate, we compared results from pollen germination and elongation obtained from TubeTracker with those made manually. We compared the manual and automatic determination of the fraction of grains that had germinated by 20 minutes of incubation in pollen germination medium. We tested each germination algorithm using pollen from six cultivars of tomato. The fractions of germinated pollen determined using the area change method were consistently lower than measurements made manually. However, none of the pairwise comparisons were significantly different from each other (Fig. 4A, purple vs. gray, p < 0.01), indicating that the automated method provides a consistent determination of time of germination. Analysis of germination using our grain to tip overlap method yielded results that were closer to measurements made manually and similarly yielded no statistical differences (Fig. 4A, black vs. gray, p < 0.01). We calculated the time of germination determined by each automated method relative to manual determination and found that the relative difference was consistent across all cultivars (Fig. 4B, each dot is a per-sample relative).

**Figure 4:**
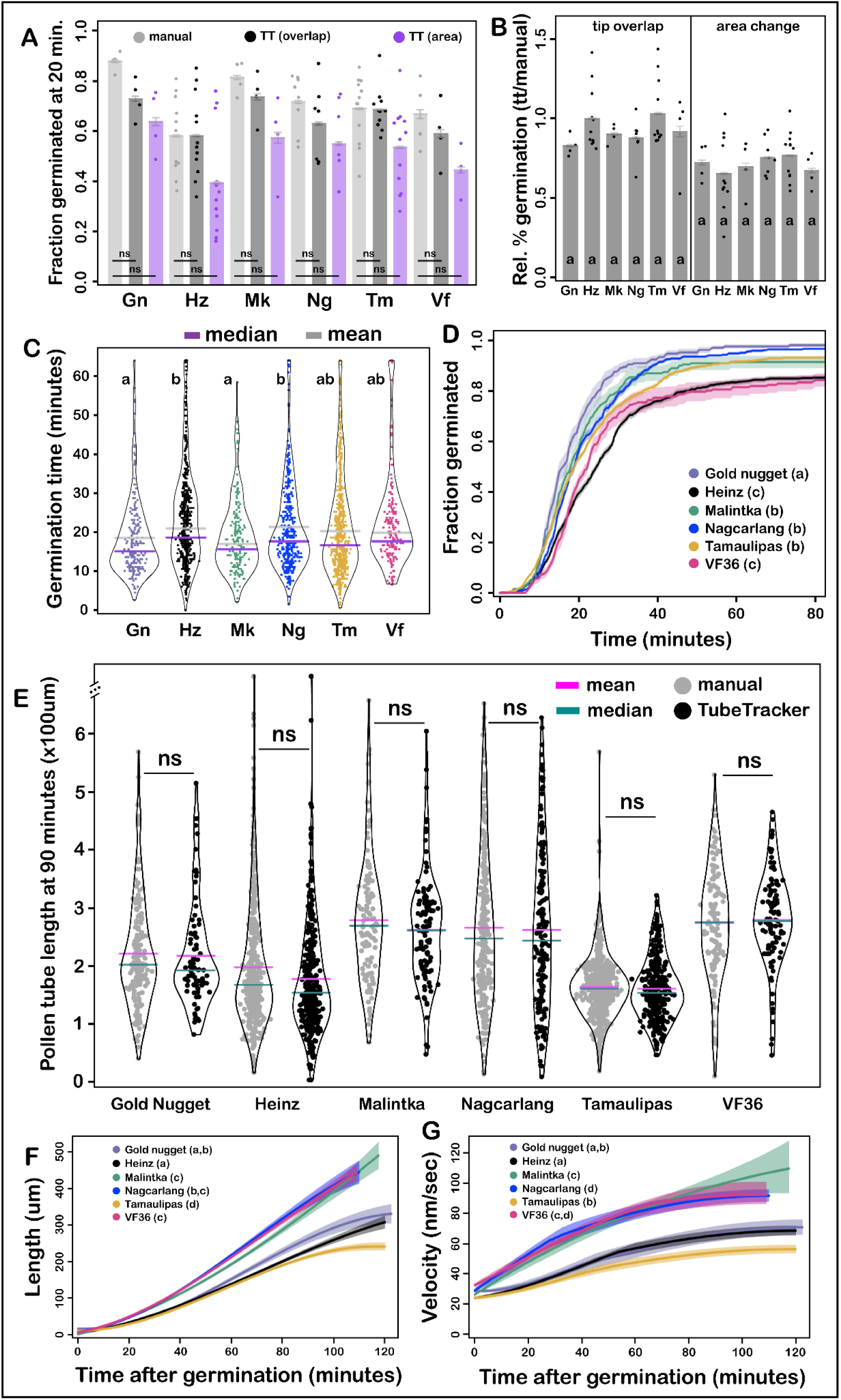
TubeTracker’s results are comparable to manual measurements and define genetic variation in tomato pollen tube performance. **(A)** Comparison of manual and TubeTracker tracked fraction of germinated grains after 20 minutes. Statistical analysis was done using the Mann-Whitney-Wilkerson U test at p < 0.01. **(B)** TubeTracker measurements of fraction germinated relative to manual measurements at the 20-minute endpoint. Each dot represents the ratio of measurements of the same sample. Statistical analysis was done as described in A. Same letters represent no statistical differences at p<0.01. The 2 different methods were not compared to each other so statistical grouping is within each tracking method. **(C)** Germination time distribution of each cultivar. Statistical analysis was done using Dunn’s test. Same letter means no statistical difference between pairs at p<0.01. **(D)** The fraction of pollen tubes germinated over time. Statistical analysis was done using survival analysis with an endpoint at 80 minutes. **(E)** Comparison between pollen tube lengths measured manually and with TubeTracker at 90 minutes. Statistical analysis was done as described in A. **(F)** Pollen tube length over time. **(G)** Pollen tube velocity over time.

We next used automated analysis to compare germination profiles across six tomato cultivars. In vitro germination of pollen tubes was found to be synchronous across all cultivars with the majority of cultivars reaching 95% germination within 30 minutes (Fig. 4C, 4D). Additionally, we found that cultivars that have been identified for high temperature fruit production (Malintka, Nagcarlang, and Tamaulipas, [2]) were also found to have higher percent germination than other cultivars (Heinz and VF36, Fig. 4D).

We then compared pollen tube length measured manually with that determined by TubeTracker. For manual measurements, the 90-minute time point was selected. For TubeTracker measurements, we recorded the longest measured length of each tube at 90 minutes of growth. Pollen tube lengths measured using TubeTracker were statistically indistinguishable from those made manually (Fig 4E). This suggests that TubeTracker can be used for endpoint analyses, but more importantly this analysis shows that the automated tool can be used to accurately and precisely quantify changes in pollen tube length and elongation rate over time.

To determine if different cultivars of tomato have pollen tubes with different elongation rates at various time points, the pollen tube length and elongation rate of six tomato cultivars were compared to one another (Fig 4F, 4G). Although all cultivars had initial elongation rates between 20 and 40 nm/sec (during the first 5 minutes), they divided into two groups composed of fast growers—Malintka, Nagcarlang, VF36—and relatively slow growers—Gold Nugget, Heinz, Tamaulipas—(Fig 4G). These differences in velocity resulted in greater pollen tube length for the fast growers compared to slow growers (Fig 4F). Finally, we discovered that after germination and throughout an initial growth period, tomato pollen tubes from Gold Nugget, Heinz, Nagcarlang, Tamaulipas, and VF-36 continuously accelerated until reaching a plateau velocity after about 45 minutes (Fig 4G). Pollen tubes from Malintka on the other end continuously accelerated and never reached a plateau velocity during our observation time.

## Discussion

The pollen tube is central to agricultural productivity and a fascinating system to study cellular extension mechanisms and their responses to biotic and abiotic stresses. However, progress has been hampered because methods to automatically measure pollen tube performance parameters using live-imaging have not been readily available. We developed TubeTracker to automate the quantification of pollen germination and pollen tube elongation, and to facilitate manual quantification of pollen tube survival (ability of pollen grains and pollen tubes to avoid cell rupture) from time-lapse brightfield images of germinating pollen grains and elongating pollen tubes. These measurements, along with their time-dependent changes, are critical to determining pollen performance under various growth conditions. We further developed TubeTracker to be interactive, user-friendly and readily implementable in a range of laboratory settings without the need for extensive training, optimization, or a prohibitively expensive imaging platform. Additionally the functionalities of TubeTracker were tested and shown to produce data useful for statistical comparisons between samples.

One limitation of TubeTracker is that the two methods it implements to automatically quantify pollen germination were found to make delayed calls of average percent germination relative to manual quantification at the specific end-point we analyzed (Fig 4A). While these effects were less pronounced for our “tip overlap” algorithm, it was still found to make delayed calls. These delayed calls were expected in cases where automated tip detection failed to detect subtle changes at the pollen aperture (detectable by an observer) as the pollen tube emerged. It was further expected that the “area change” algorithm would return rates lower than fully manual quantification because its reliance on the increase in box size past a user-defined threshold will likely miscall germinated pollen grains for which the bounding box remains below that threshold (Fig. 3C). Despite this limitation, the uniformity of error across all experiments suggests that using either of the automated methods would greatly facilitate making comparisons between genotypes and/or treatments as long as all these results were generated using the same germination method and user-defined parameters.

The ability to track pollen survival over time (defined as the fraction of grains and/or tubes that avoid cell rupture over time) is critical to determining pollen performance. To automate this analysis, we would need to develop an algorithm that classifies pollen tube rupture. Unfortunately, we have not been successful using available approaches or using analogous algebraic and geometric principles such as those used for grain and tip detection (circularity of grains and edge of the line segment produced by the skeleton of a pollen tube respectively, Fig. 2C, 2D). However, TubeTracker facilitates manual tracking of cell rupture events by providing easy to follow instructions (Sup. File S1) and by incorporating past detected events into the quantification of survival rates in later frames. Importantly, TubeTracker provides a means to produce training data sets that can be used to implement and improve classification approaches that take advantage of machine learning.

While TubeTracker allows for the extraction of key pollen grain and tube performance parameters, further improvement can be achieved to increase its usefulness and obtain a more complete view of pollen performance. For example, while it can track the displacement of a tip, TubeTracker is currently incapable of measuring pollen tube tip thickness or bulging which are important phenotypes that can arise due to stress (Sup. Fig. S2). Future releases of TubeTracker will take advantage of its excellent “segment edge tip detection” method to locate tips and calculate the fraction of the bounding box area covered by positive tip pixels over time. This fraction will then be used to estimate the relative thickness of each tip and how it changes over time. TubeTracker uses separate sets of approaches to define pollen grain germination, tube elongation, and pollen tube rupture (Figs. 2 and 3). Consequently, these three parameters cannot be linked together which limits our ability to answer interesting questions like whether the tip accelerates or slows down right before it bursts. Future releases will incorporate the ability to assign an elongating tip to their grain of origin given proper quantification. This will be done by developing a method that searches for overlaps between the location of each pollen grain and the location at which each pollen tube track starts.

TubeTracker will allow our field to address key questions about how pollen grains and pollen tubes respond to a variety of perturbations using richly quantitative data that track key features of pollen performance over time. For example, how does germination at high temperature affect the fitness of pollen tubes from the different tomato cultivars we have explored so far? Additionally, do thermotolerant cultivars, which have been shown to have improved fruit and seed production due to the maintenance of high temperature pollen tube growth in the pistil [2], have pollen with improved grain germination, tip elongation and pollen cell integrity at high temperatures compared to thermosensitive cultivars? These experiments will allow us to make associations between different cultivars and specific features of pollen performance that require improvement. TubeTracker will also be very useful in analysis of specific mutations that affect pollen performance. For instance, it is proposed that the main ROS producers in pollen tubes are RBOH H and J. *rbohh*/*rbohj* double mutants resulted in pollen tube burst under optimal conditions [42–44]. Using TubeTracker, we can define the specific pollen performance parameters that are regulated by these key genes and determine whether inhibition or super activation of RBOH enzymes will improve pollen performance. TubeTracker will further allow us to define how application of abiotic stresses (e.g. salt/osmotic stress) or small molecules (e.g. hormones and regulatory proteins like Rapid Alkalinization Factors, RALFs) regulate each aspect of pollen tube germination, elongation and survival. Finally, although we developed TubeTracker to be used for tracking pollen, biologists studying a range of tip-growing cells (e.g. root hairs, neurons, and fungal hyphae) can also use this tool to extract quantitative data over time.

## Materials and Methods

### Tomato Cultivars, plant growth, and pollen growth medium

Seeds of 6 tomato cultivars: Heinz 1706 - BG (LA4345), VF36, Tamaulipas (LA1994), Malintka 101 (LA3120), Nagcarlang (LA2661) and Gold Nugget (LA4355) were obtained and maintained at Brown University as described previously [2]. The growth medium used for in-vitro live imaging was prepared as explained previously [2]. For each experimental run (1 in late fall 2021 and 2 in summer 2023), a masterbatch for media was prepared and stored in 10 ml aliquots at -20°C.

### Live Imaging System (Sup. Fig. S3)

To acquire the time-lapse series used to develop and test TubeTracker, we set up an imaging system that efficiently recorded pollen tubes growing on a 2D surface. Each time-lapse series was made using a Swift 5.0 Megapixel Digital Camera mounted on the eyepiece of an OMAX 40X-2000X LED Binocular Compound Lab Microscope. An AmScope TCS-100 Microscope Temperature Control Stage Slide Warmer was fixed on the microscope stage and a secondary petri dish filled with water was placed on it. The whole system was placed inside a Cole-Parmer H2200-H-C-E Mini Digital Incubator and covered with plastic to protect the system from incubator air currents. Closed Glass bottles filled with water were also added to the incubator to help stabilize the ambient temperature (absorb extra heat or release heat after the incubator was opened) and an EasyLog temperature logger temperature probe (EL-USB-TC-LCD) was permanently kept in the water-filled secondary container to accurately record its temperature which was also assumed to be that of the sample. The camera was connected to a computer on which the Swift software was installed. All recordings were done at low light intensities on the microscope. Two of these systems were created to allow for parallel recordings.

### Movie Acquisition

Pollen was collected from 4 to 5 flowers as described in [2] using a modified vibrating toothbrush, collecting into a 0.5 ml Eppendorf tube less than 30 minutes before each experiment. An aliquot of the collected pollen was then thoroughly mixed with 1 ml PGM pre-warmed to 28°C and added to a 35x10 mm clear petri dish. The dish was then placed inside the secondary container of the imaging system and allowed to incubate for 3 minutes to allow pollen grains to settle down at the bottom. After 3 minutes, an additional 3 ml prewarmed PGM was slowly added to the side of the imaging plate. Once a proper field of view was selected, the movie was recorded for at least 125 minutes at a rate of 14 frames per second. Less than 10 minutes elapsed each time between the addition of pollen grains into PGM to the start of recording. After the recordings were completed, the movies were then reduced to 1 frame every 30 seconds. To increase contrast of these (not necessary for TubeTracker), the frame selected every 30 seconds was the average signal from 2.5 seconds prior and after the set time point.

### Data quantification

Manual quantification of germination was done by hand counting how many grains had properly germinated at the endpoint of each movie (20 minutes). Manual quantification of pollen tube length was done by manually tracing the path of each pollen tube at the 90 minute endpoint using ImageJ. All automatic measurements were made using automated and manual functionalities of TubeTracker (https://github.com/souonkap/TubeTracker).

### Data analysis and visualization

to process the data after quantification, we developed code in R that uses the raw quantification files to perform different types of analysis and generate phenotypic profiles such as those shown in this paper. Statistical analysis of this data was done using survival analysis [22–25] for germination profiles, burst profiles, and grain response, or using a combination of ANOVA and Kruskal-Wallis tests to test for significant variation in the data. Once variation was detected, a combination of the Mann-Whitney-Wilkerson U test, Welch two sample t-tests, and Dunn’s test were used to determine statistical significance in pairwise comparisons. Details on which test was used for each experiment can be found in figure legends.

### Github repository and user feedback

TubeTracker is a tool developed by a biologist with a passion for object-oriented programming. We encourage the community to report bugs and provide their suggestions for improvement. We further encourage users to independently improve upon our tool and have provided the complete python code at https://github.com/souonkap/TubeTracker, along with installation instructions and a video sample for training purposes.

## Acknowledgements

This project was funded by a grant from the US NSF (IOS-1939255, Co-PIs MAJ, RP, GKM, AL) with additional support from a USDA/NIFA grant (2020-67013-30907, Co-PIs GKM, MAJ) and Brown University Presidential Fellowship (S.Y.O.). We thank Sherry Warner (Brown University) and Nick Vasquez (Brown University) for their expertise in growing and maintaining tomato plants. We thank our former colleague, Dr. Nathaniel Ponvert for sharing his insight on the development of an earlier tracking algorithm called ASIST. We thank former Brown University undergraduate Kelly Pan for pointing S.Y.O. toward the training material used to start the development of this software. We further thank Dr. Alison DeLong for her feedback on the manuscript as an outside reader.

## Author contributions

S.Y.O. designed the algorithms, wrote the scripts, and analyzed data;

S.Y.O. M.J. designed the experiments;

S.Y.O., B.O., Y.O., tested all Python algorithms and functionalities and quantified the data;

S.Y.O. generated the figures and illustrations

S.Y.O. M.J. wrote the manuscript with input from B.O. and Y.O..

## Supplementary Materials

**Supplement Figure 1:**
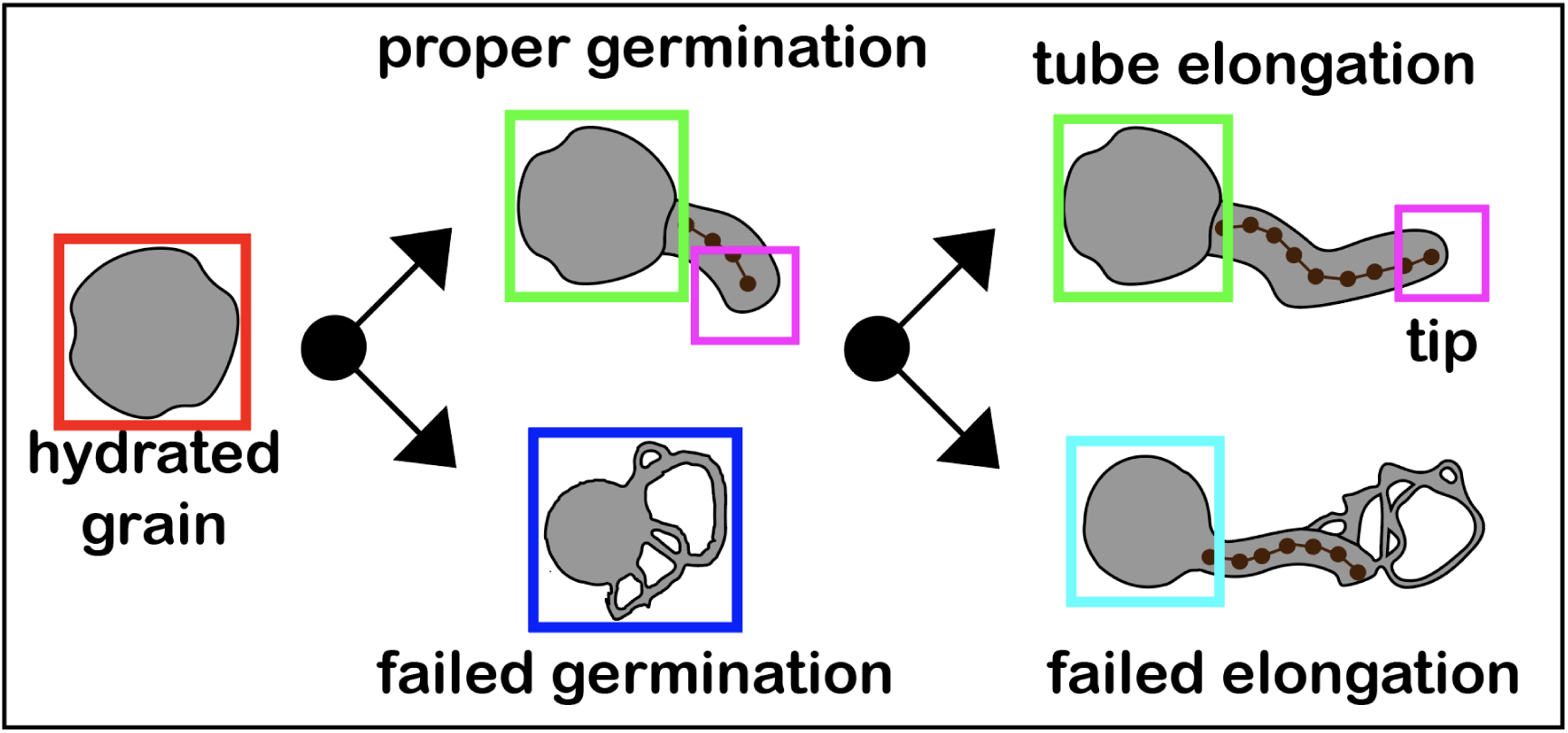
Different features that can be extracted from pollen for survival and elongation tracking using TubeTracker.

**Supplement Figure 2:**
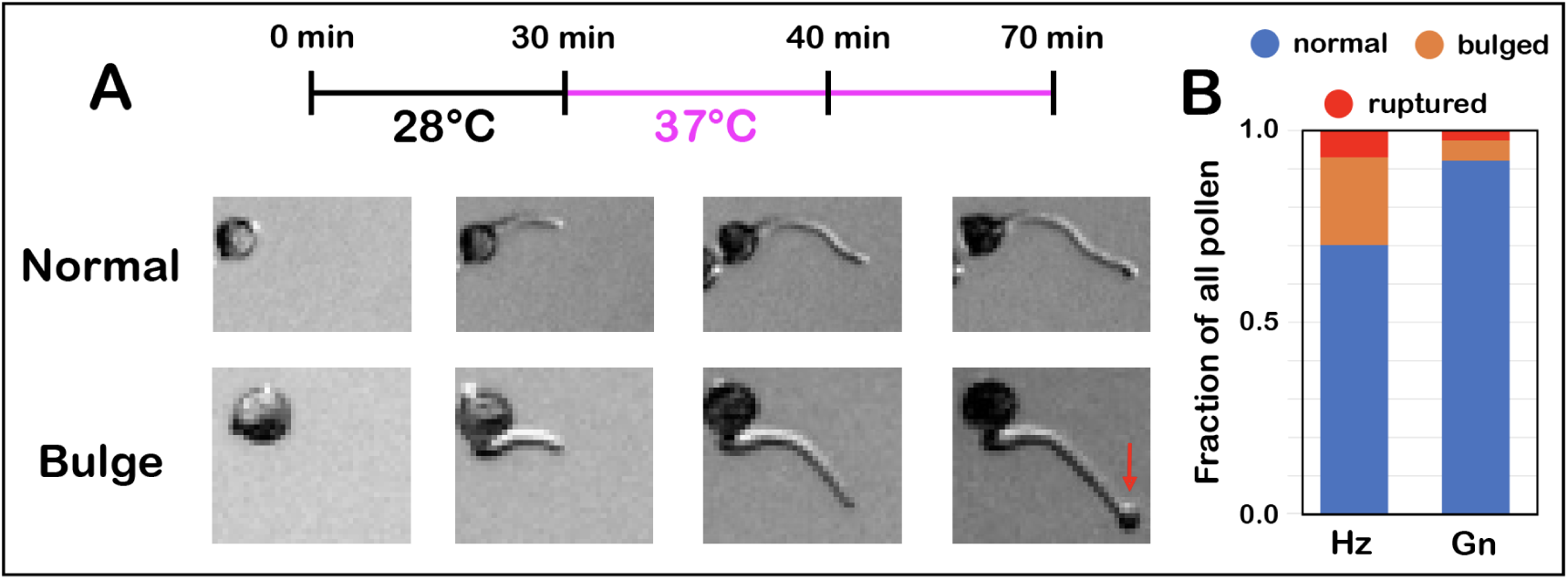
High temperature on germinated pollen tubes causes pollen tube tip bulging (swelling). (**A)** Live imaging experimental conditions and representative images of bulged and normal pollen tubes. (**B)** Quantification of bulging of pollen tubes from Heinz (Hz) and Gold Nugget (Gn).

**Supplement Figure 3:**
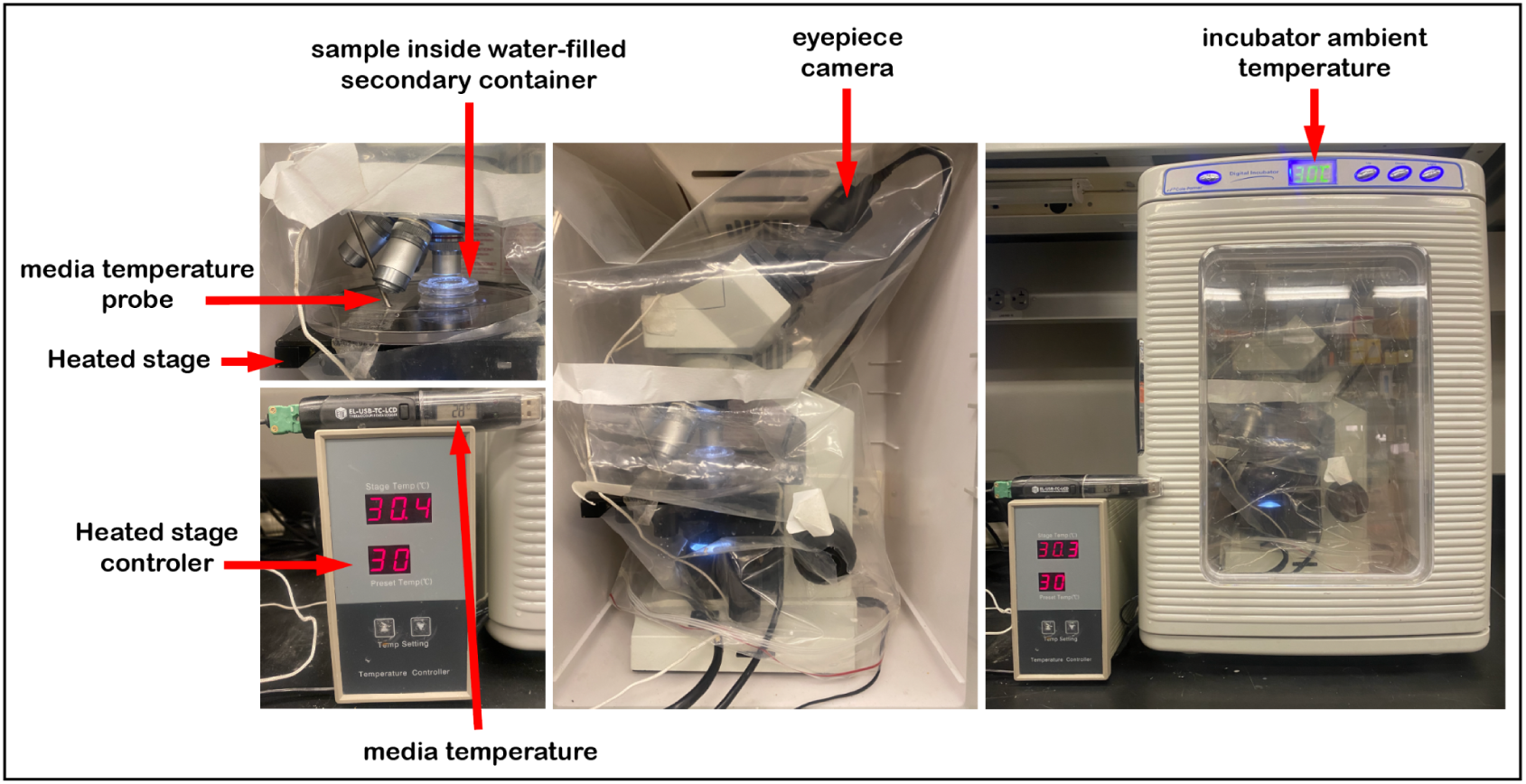
Live Imaging system used for movie acquisition.

