## Supplemental File S1 for "Semi-Automated High Content Analysis of Pollen Performance Using TubeTracker"

### TubeTracker Manual

Sorel V Yimga Ouonkap
Johnson Laboratory
Brown University

### Installation Instructions

### Install a virtual environment handler (miniconda3)

We recommend installing TubeTracker in a virtual environment to isolate it from other programs on your system.

Miniconda3 is commonly used to do this. Installing miniconda will add the command “conda” on which the installation and launch scripts of this program depend on.

To install the command, please visit the site bellow and download the miniconda3 version compatible with your Mac or other apple computer. Please download only the “pkg” versions of “Minoconda3 macOS Apple M1” or “Minoconda3 macOS intel”. After download, launch the install and follow the prompt.

Link: https://docs.anaconda.com/miniconda/

### Installing TubeTracker

After installing the conda command, double click on the provided file called "Install_TubeTracker" to start the installation of the application dependencies and follow the prompt.

You will mainly be asked to confirm the installation of certain dependencies. Enter "y" or "yes" then press "Enter" to continue.

Do not worry about giving permission. As stated before, we create a virtual environment to install these packages and they are isolated from everything else on your computer. They will not affect any other programs you currently have installed on your laptop.

If the installation completes without any issues, the tool should be ready to launch and you can launch it using "Start_TubeTracker". If the installation was successful, the graphical user interface of TubeTracker should appear

### Installing TubeTracker

These dependencies will be installed

- Python (version 3.11.4)
- Wxpython (version 4.2.1)
- Opencv (version 4.10.0)
- Pandas (version 2.1.4)
- Motpy (version 0.0.10)

### Usage Instructions

### TubeTracker GUI

Loading and segmenting movie + saving results

Manual tracking and quality control

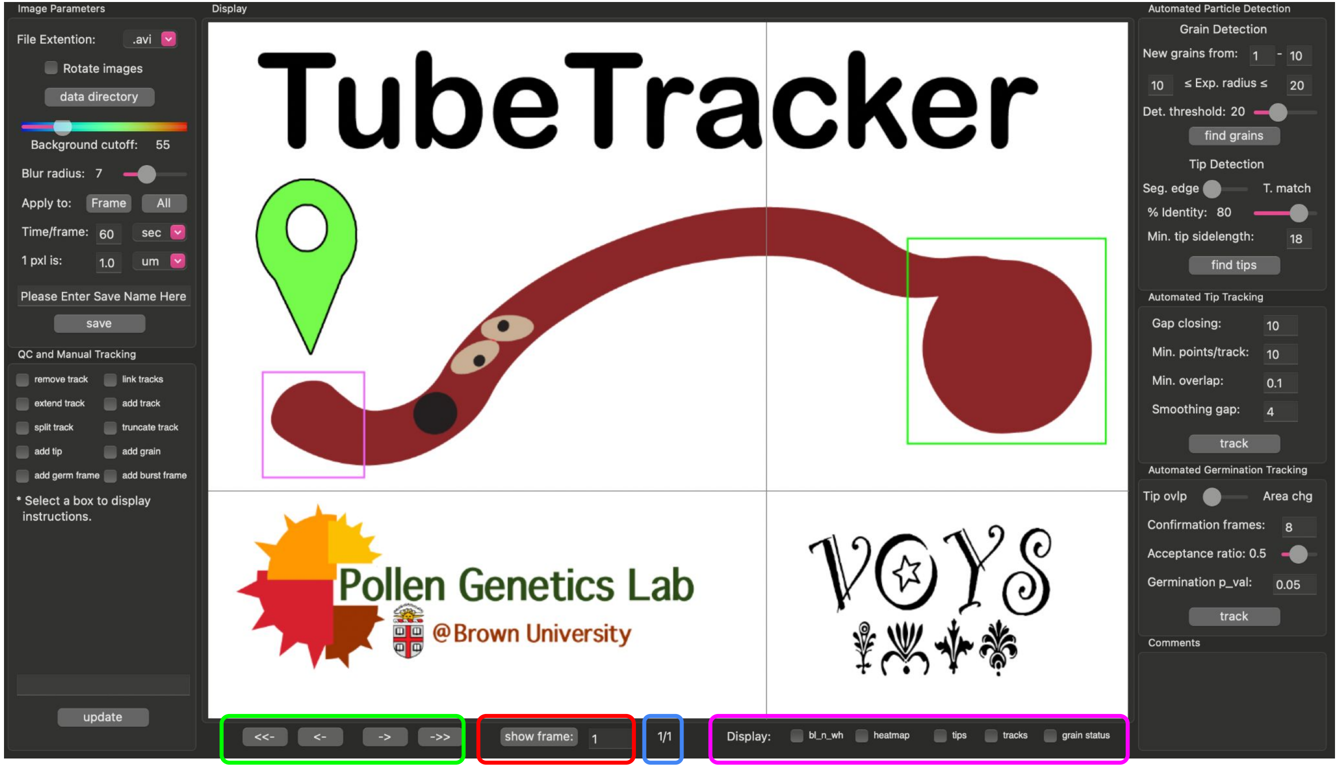


Automated grain and tip detection

Automated tracking of grain germination and tip elongation

Important comments
Currently inactive

| Move 1 or 5 frame(s) backward (**<-**, **<--**) or forward (**->**, **-->**) respectively. | Show selected frame | Current frame number / total frame number | Display various data types when available |
|---|---|---|---|

**Screen Controls**

### Usage Instructions

#### Loading a time-lapse series

### Movie quality and dimensions

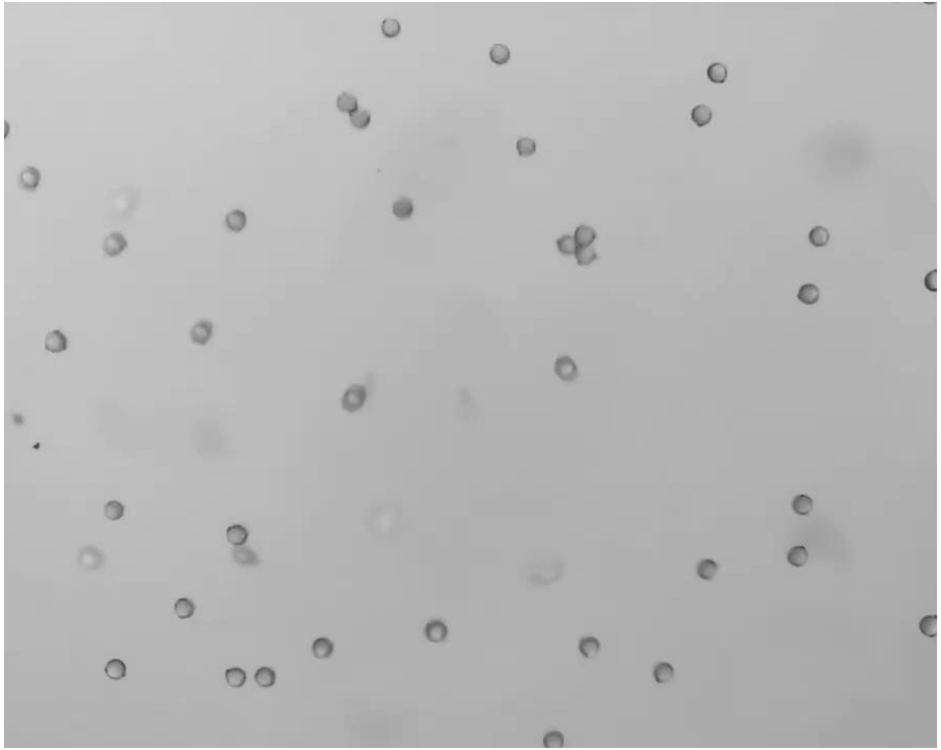

2 hour movie, 1 frame every 60 seconds

- Works with brightfield or DIC movies or series of images
- Avoid high intensity background as much as possible. The movie shown provides an acceptable level of background
- Currently, any loaded movie will be resized to 1000/755 w/h. This may distort the movie however, the distortion is remembered and accounted for during data quantification.
- Movies can be rotated to minimize distortion
- Avoid unwanted motions such as lateral grain and tube displacement.

### Loading and segmenting a movie into TubeTracker

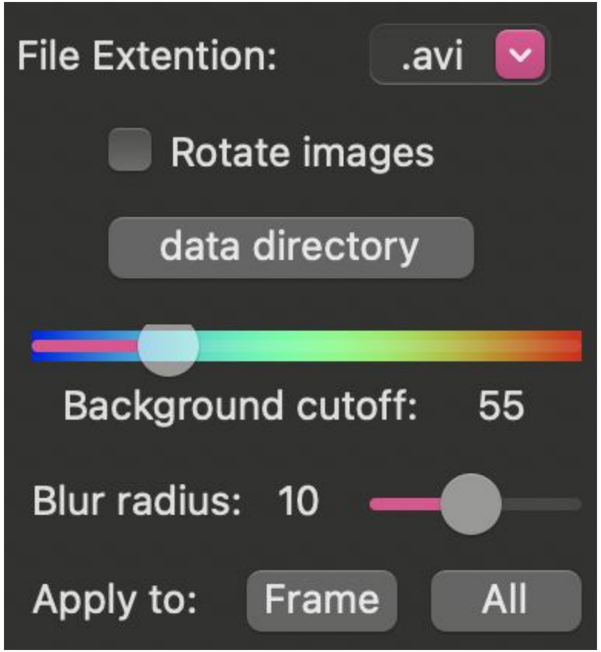


- 4 pieces of user-provided information are needed to load and segment a movie.
- 2 are required for loading a movie
  - The expected **"File Extension"** (next slide)
  - **"Rotate Images"**: Optional, rotate the movie 90° counterclockwise.
- 2 are required to differentiate foreground from background
  - **"Background cutoff"**: the pixel intensity used to remove background. All pixel intensities below this threshold are considered background
  - **"Blur radius"**: used to alter each movie to make edges of pollen grains/tubes more uniform (see slide 14)

### Loading a movie into TubeTracker

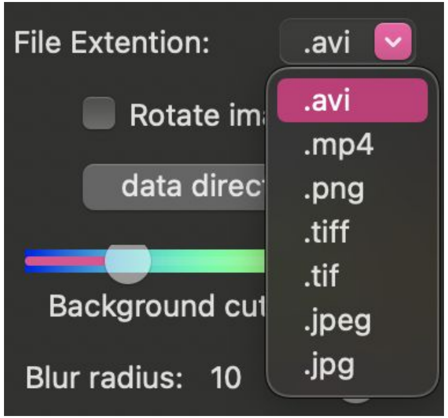


- When Loading a movie, First select its file extension.
- TubeTracker supports 7 file formats.
  - 2 movie formats: "**.avi**", "**.mp4**"
    - If one of these is chosen, you will be expected to select a single movie file with the chosen format
  - 5 single image formats: "**.png**", "**.tiff**", "**.tif**", "**.jpeg**", "**.jpg**"
    - If on of these is chosen, you will be expected to select a folder holding a series of images with the chosen format are stored.
    - **NB: the temporal arrangement of these single images is done based on the alphanumeric order of filenames in the chosen folder (a to z; first to last).**
- Once the file format is selected, click on "data directory" to select a movie or folder
- When loaded, the movie frames will be displayed automatically

### TubeTracker Screen Controls

#### Navigating the time-lapse

Move 1 or 5 frame(s) backward (**<-, <--**) or forward (**->, -->**) respectively.

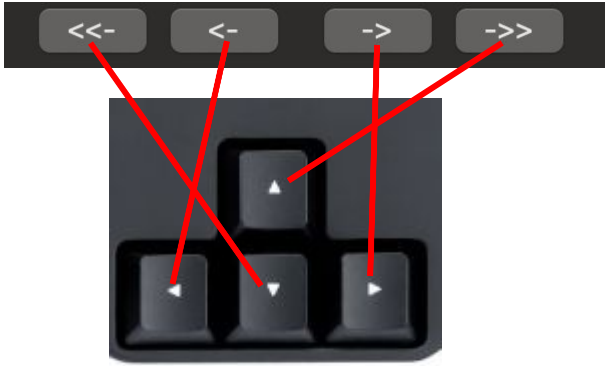


**Keyboard shortcuts:**
If unresponsive, move the cursor on the screen display.

Displaying a specific frame

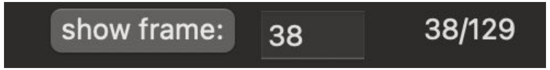


#### Displaying various image/data info

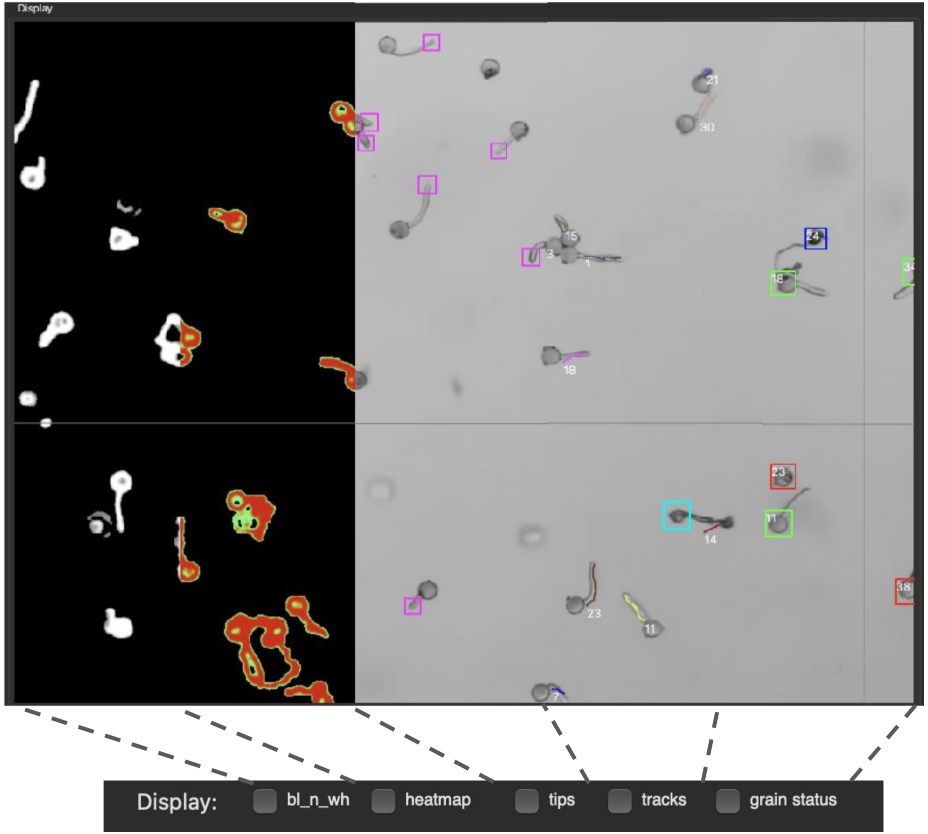


grain status:
Detected /
Ungerminated
Grain burst
Pollen Tube burst
Germinated

### Movie segmentation

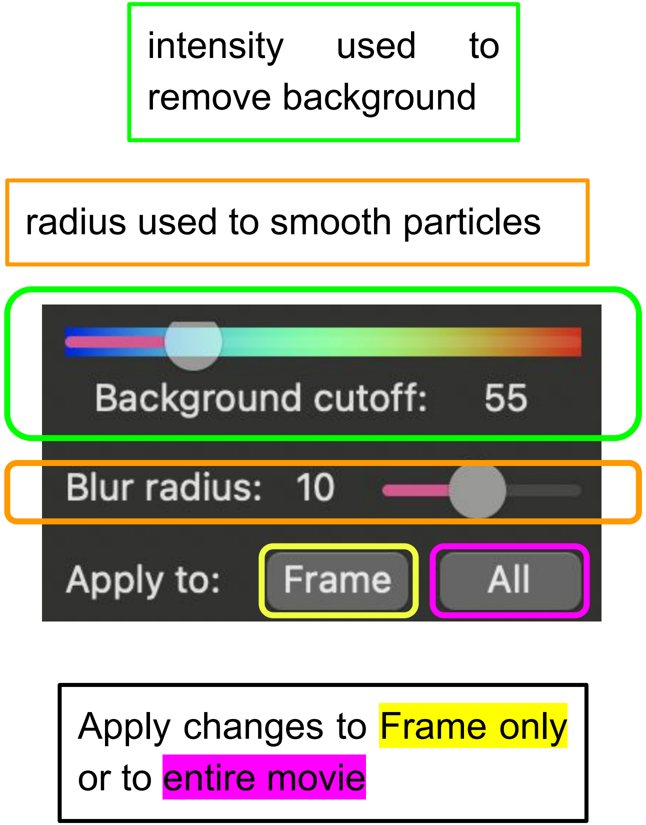


- A main issue with video analysis is the difference in the camera used to collect images. For machine learning applications for example, the algorithms often underperform when the collection camera is changed. To avoid these camera issues and make the tool readily useable, TubeTracker first differentiates foreground from background (segmentation) to produce black and white images for which particle edges are highlighted based on local contrast differences.
- The black and white images (which no longer depend on camera specs) are then used for downstream applications.
- TubeTracker does most of the segmentation without user input. However, with high sensitivity to contrast changes, some low intensity noise can still be present and some pollen may not be detected.
- To resolve these issues, it is necessary to adjust the intensity to use for background subtraction and the blur radius that will be used to make particles and edges more uniform.
- See next slides for examples

### Adjusting the background cutoff

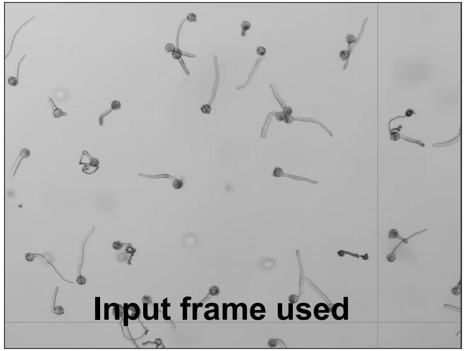


To select the correct background cutoff, you can use the heatmap (slider, above) to guide your selection.

It is possible to display color coded pixel intensity values by selecting **"heatmap"** under **Screen Controls** at the bottom of the GUI.

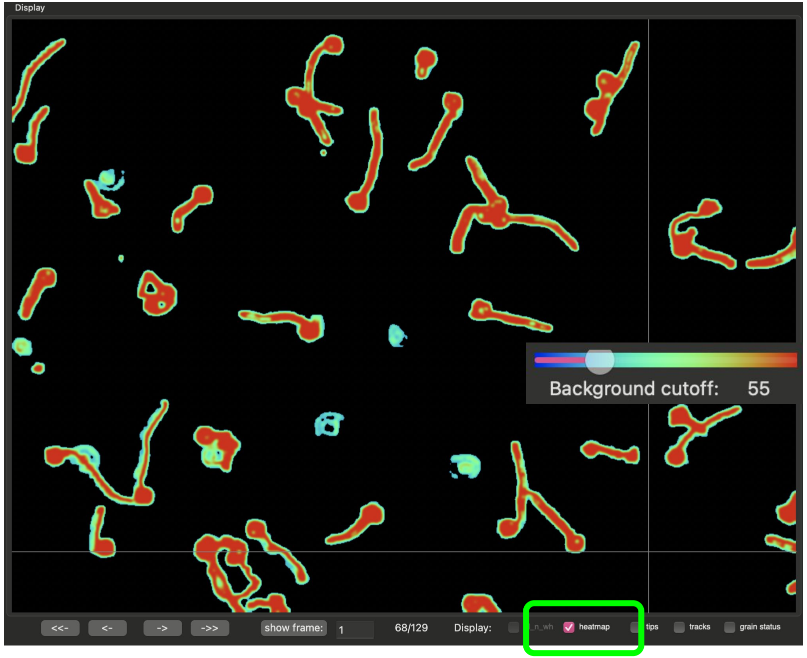


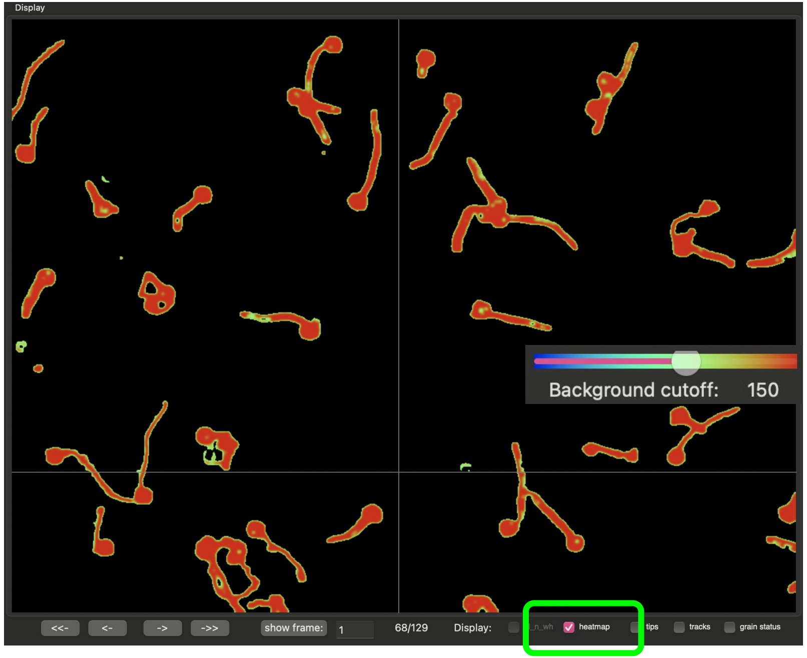

### Usage Instructions

#### Automated and Manual Particle Detection

### GUI panel used for detecting particles

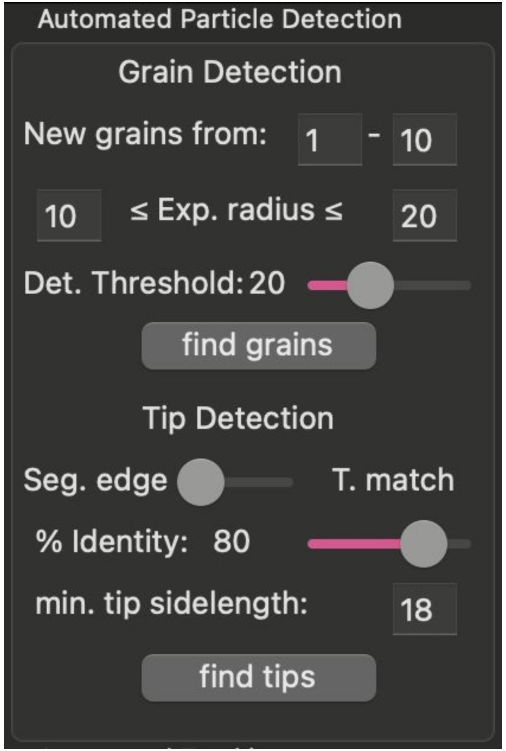


Once a movie is loaded and segmented, the user can use manual and automated options to detect target particles such as pollen grains and pollen tube tips. Here, I describe how to detect these two particle types using an automatic approach that can be modified manually.

### Automated grain detection

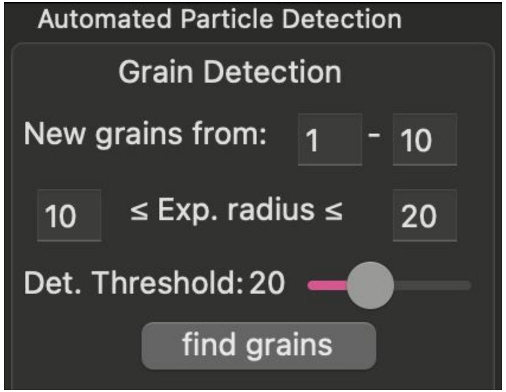


- Grains in TubeTracker are assumed to be circular particles and this property is used to locate them in space and across time.
- Once grains are spatially identified in each frame, they are temporally tracked to update their location over time. If a grain is temporally lost, the location of its last recorded position is used in subsequent frames.
- 3 pieces of user-provided information are needed to detect grains.
  - **“New grains from:”**: the range of frames used to locate ungerminated grains for analysis (grains not detected in these frames will be ignored)
  - **“Exp. radius”**: the expected range for the radius of grains (pixels). To choose an appropriate range of values, you’ll need to “add” and then “remove” a few grains manually (see next slide).
  - **“Det. Threshold:”**: a threshold value used to determine if particles meet expectations for circularity.

### Automated grain detection, example

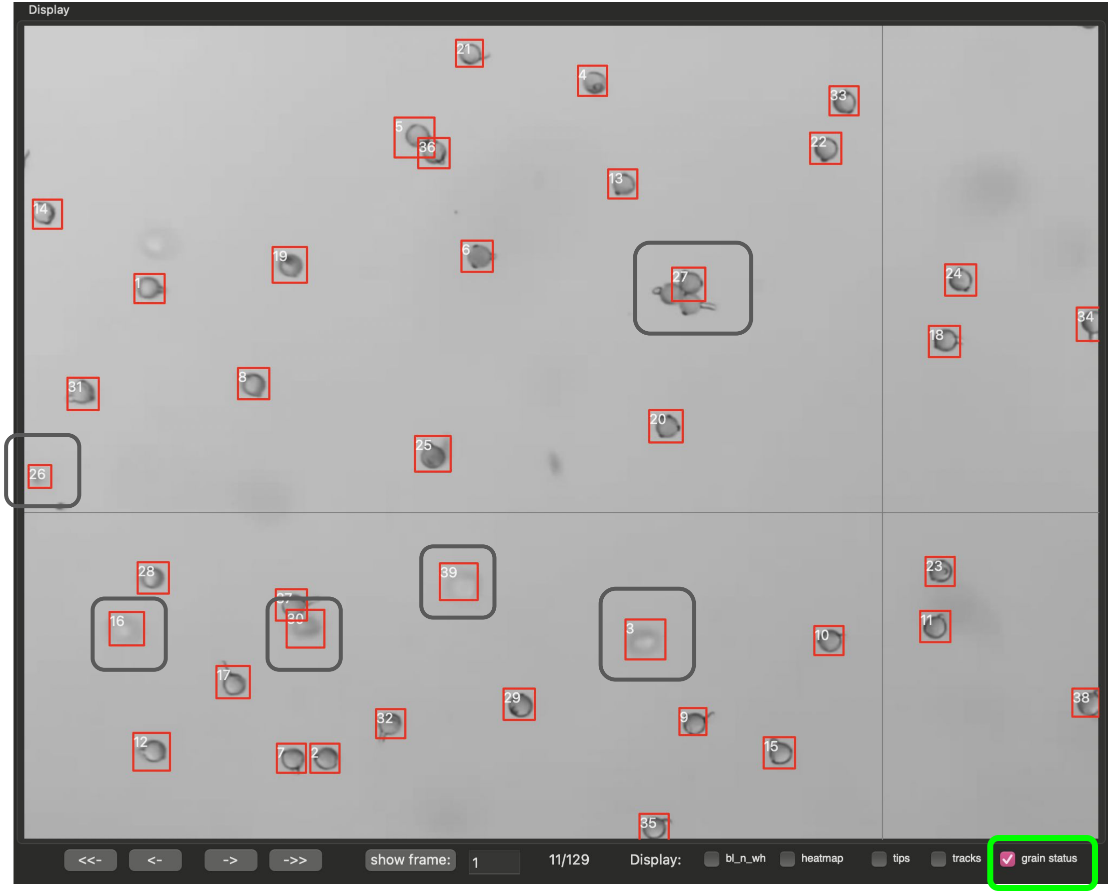


Detected grains can be displayed using the **“grain status”** option of **Screen Controls**

Note: When Det. Threshold was set to 17 and defaults were used for all other parameters, all but 2 grains were detected. These undetected grains however were in a clump (such as those close to grain 37).

Further note that 5 particles were falsely identified as grains (grains 3, 16, 26, 30, 39) because they were noise that resembled grains. These grains need to be removed using manual options

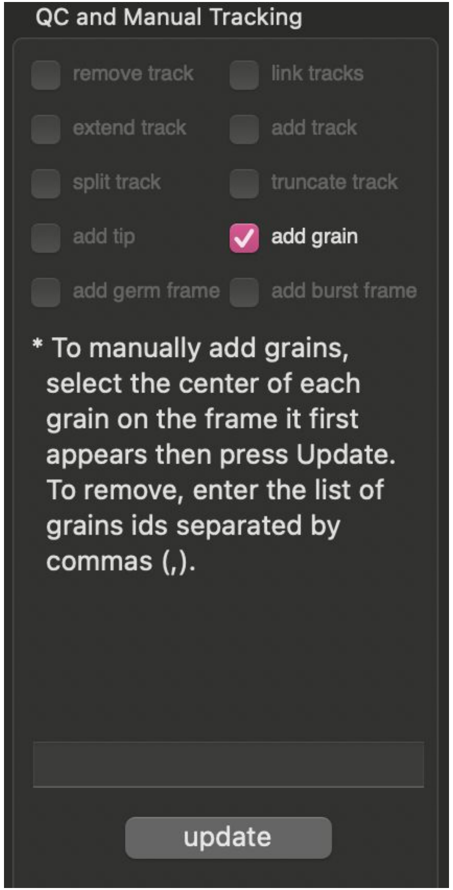


### Manual grain detection

- Grains can be added and removed using the **“add grain”** option of the QC and Manual Tracking panel.
- Follow the instructions provided in the panel after selecting needed feature (e.g. “add grain”)
- Using this tool, the false positives from the previous frame (grains 19, 24, 30, 36) were removed and the missed grains were detected. The grain ID #s for removed grains are re-assigned to any user defined grains.

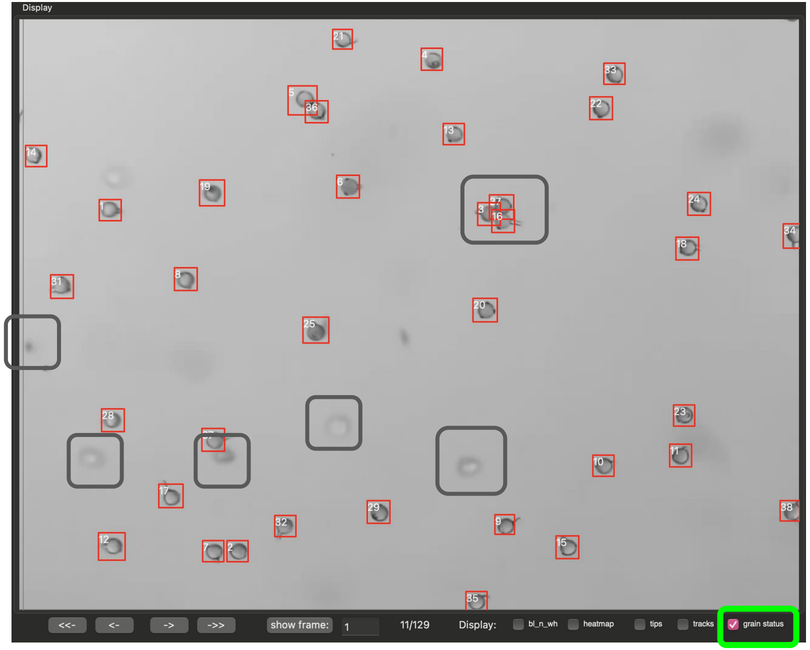

### Automated Tip detection

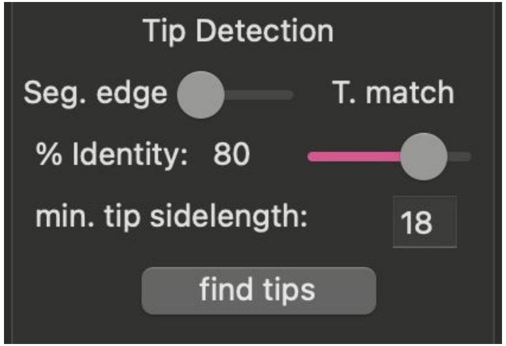


TubeTracker can implement either of 2 tip-detection algorithms - use the provided toggle to indicate which you are using.

- **“Seg. edge”**: (segment edge) this is the default and preferred algorithm because of its high detection rate when images are properly segmented and fast processing time. Only the **“min. tip sidelength:”** (in pixels) parameter is required for this algorithm.
- **“T. match”**: (Template Match) this algorithm works by searching for instances of predefined tip images based on a percent identity. This algorithm is time consuming especially with large number of frames but works well when segmentation is not optimal. This algorithm requires the **“min. tip sidelength:”** parameter as well as the **“% Identity:”** parameter which determines the degree to which a newly detected tip must resemble a predefined one.

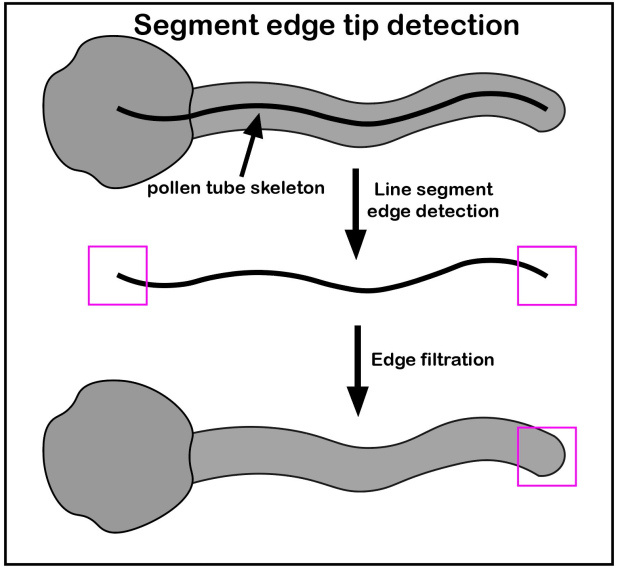


**Template matching tip detection**

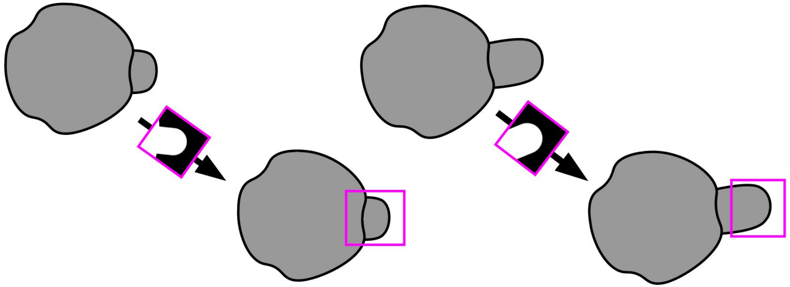

### Automated tip detection, example

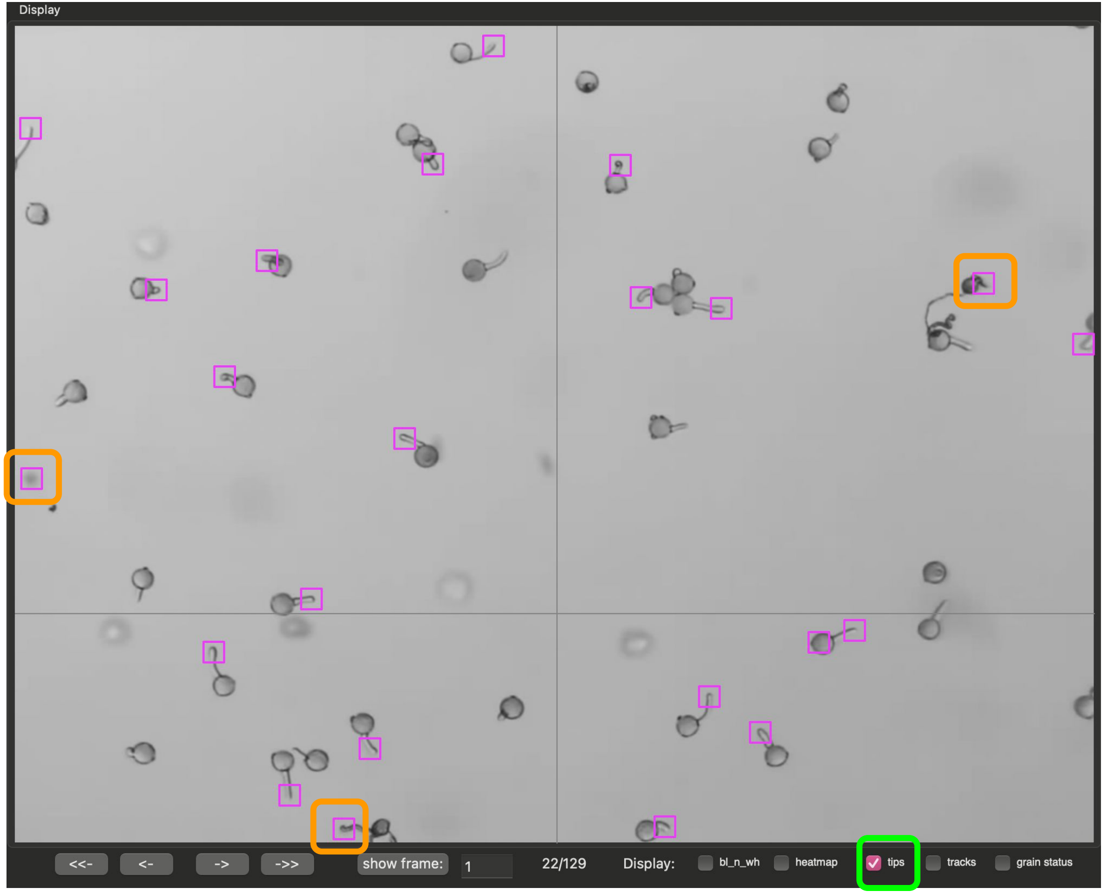


Using the **“Seg. edge”** algorithm, the majority of the tips can be detected.

In this case, we’re viewing frame 20.

It is okay if some tips are skipped because missing tips can be added manually. Additionally, automated tracking of germination and tip elongation can accommodate tips missing in some frames as long as it does not persist over a large number of frames.

Please note that some noise and dead grains spills were detected as a tip but accounts for a small portion of all tips. These can be manually removed.

Further note that tips can be displayed using the **“tips”** option of Screen Controls

### Manual tip detection and modification

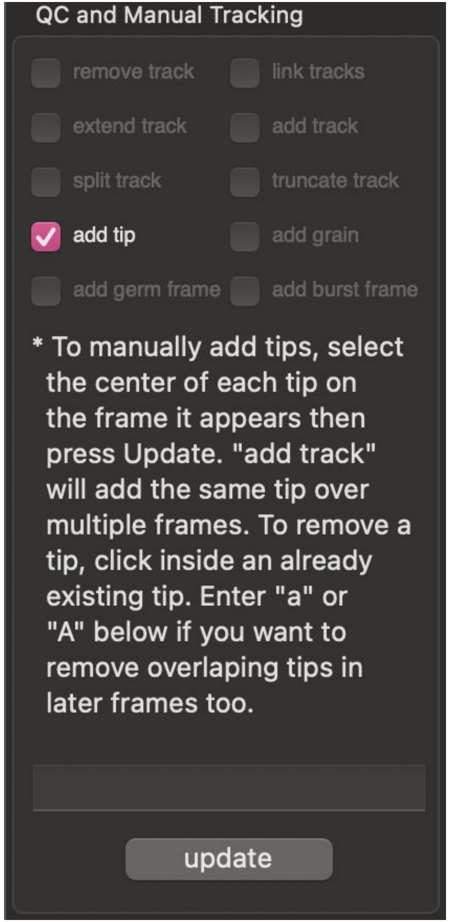


- Tips can be added or removed using the **“add tip”** option of the “QC and Manual Tracking” panel.
- To manually add or remove a tip, follow the instructions provided.
- Note in the frame below that falsely identified tips were removed while others have been added

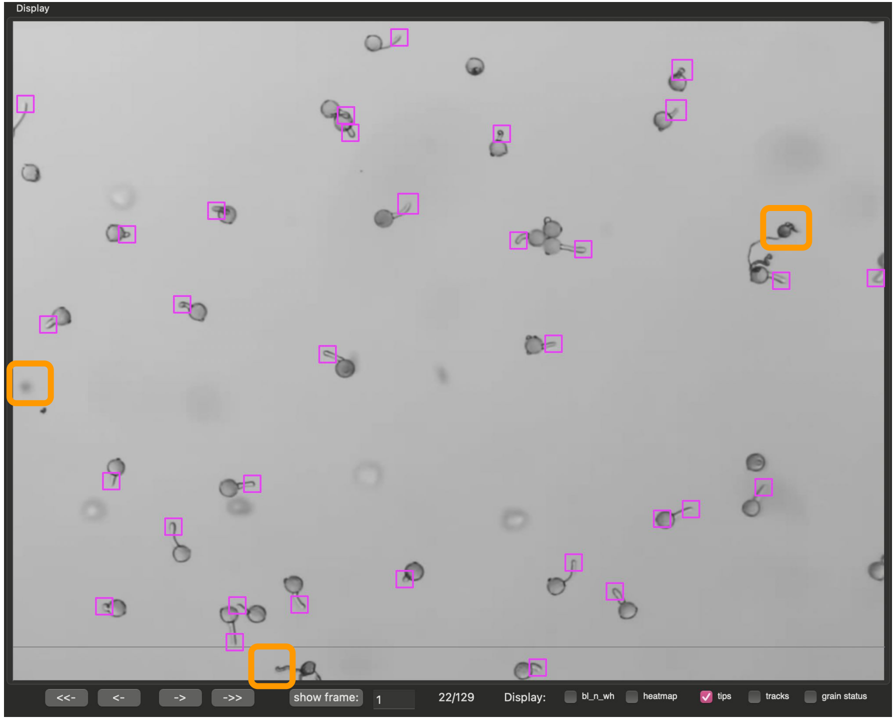

### Usage Instructions

#### Automated and Manual Tracking of Tip Elongation

### Automated tip tracking

TubeTracker implements the multi-object tracker from motpy to connect overlapping tips in successive frames.

To track tip elongation, 4 parameters are needed:

- **“Gap closing”**: This is the number of successive frames over which a tip is allowed to disappear
- **“Min. points/track”**: This represents the minimum number of frames over which a tip must be detected to yield a valid track
- **“Min. overlap”**: This sets the minimum overlap (ratio of the intersection to total area) allowed between the bounding boxes defining tips in successive frames. If this is exceeded and gap closing fails, a new track will result rather than forming a continuous track. These tracks can manually be connected to form a continuous track (**see tracks 3, 43, and 94 below**).
- **“Flattening gap”**: this represents the resolution used for producing less wrinkled tracks. It is measured in number of frames and cannot be greater or equal than **“Min. points/track”.** See example below.

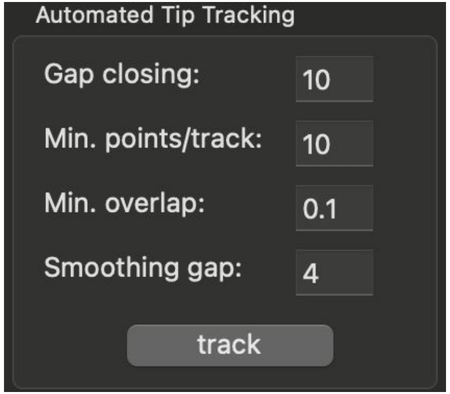


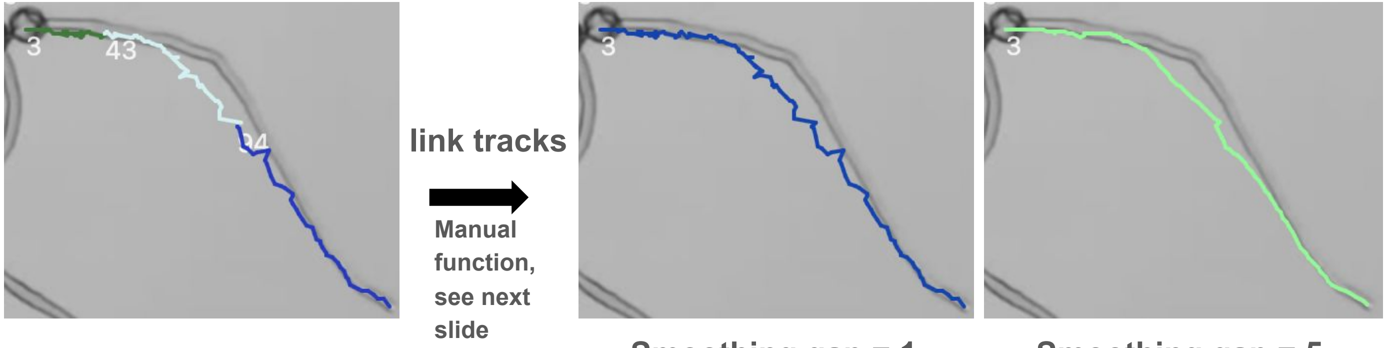


Smoothing gap = 1

Smoothing gap = 5

### Manual user inputs improve automatically generated tracks

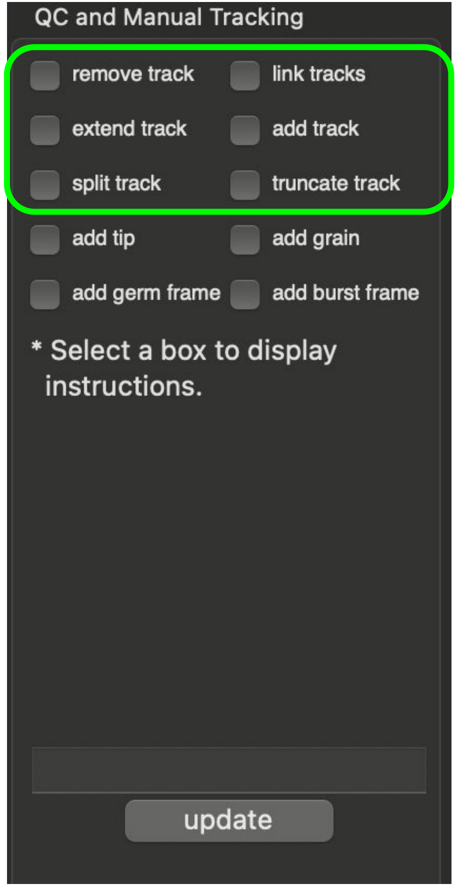


Pollen tube tracks generated through automated functions can be modified using manual functions highlighted inside the green box. Note that instructions for how to modify tracks are provided when each option is selected.

Six options are provided to manually manipulate each track directly on the screen or by indexing tracks using their track IDs.

These options are:

- **“add track”**: creates a new track using user provided screen positions over multiple frames.
- **“remove track”**: removes one more more existing tracks given their ids
- **“link track”**: links together a series of tracks belonging to the same pollen tube given their ids.
- **“extend track”**: extends an existing track using user provided screen position over one or more frames.
- **“split track”**: cuts an existing track into 2 new tracks given the former track id and a screen position corresponding to the split location
- **“truncate track”**: similar to split, however, the tail end of the former track is discarded.

### Usage Instructions

#### Automated and Manual Tracking of grain germination and grain/tip survival

### Automated tracking of grain germination

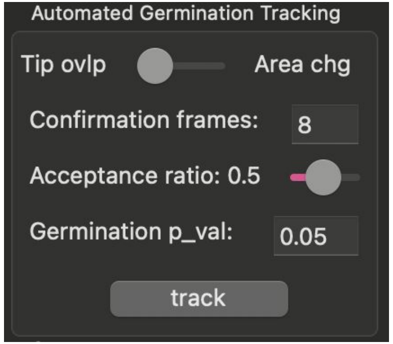


TubeTracker implements 2 algorithms for tracking germination. Use the provided toggle to indicate which you are using.

- "**Tip ovlp**": This method checks for overlap between detected grains and tips to make germination calls and is the preferred method in cases where automatic/manual tip detection is complete across all/most frames.
- "**Area Chg**": This methods uses the increase in area of the bounding box around the germinated grain to make germination calls and is the preferred method when tip detection is inconsistent or was not done across frames.

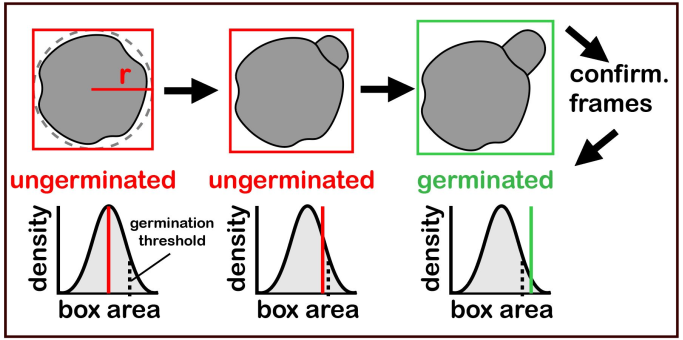


3 user inputs are required to track germination

- "**Confirmation frames**": number of frames used to confirm that a tube has emerged from a grain.
- "**Acceptance ratio**": the fraction of "**Confirmation frames**" for which a grain must remain germinated during confirmation.
- "**germination p_val**": threshold from null distribution used to make germination calls. Only required when using the "**Area Chg**" algorithm.

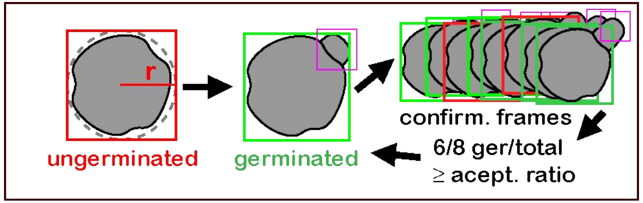


Panel B (tip overlap) and C (area change) of Figure 3 of the manuscript of TubeTracker demonstrating the algorithms used for tracking elongation.

### Automated germination tracking, an example

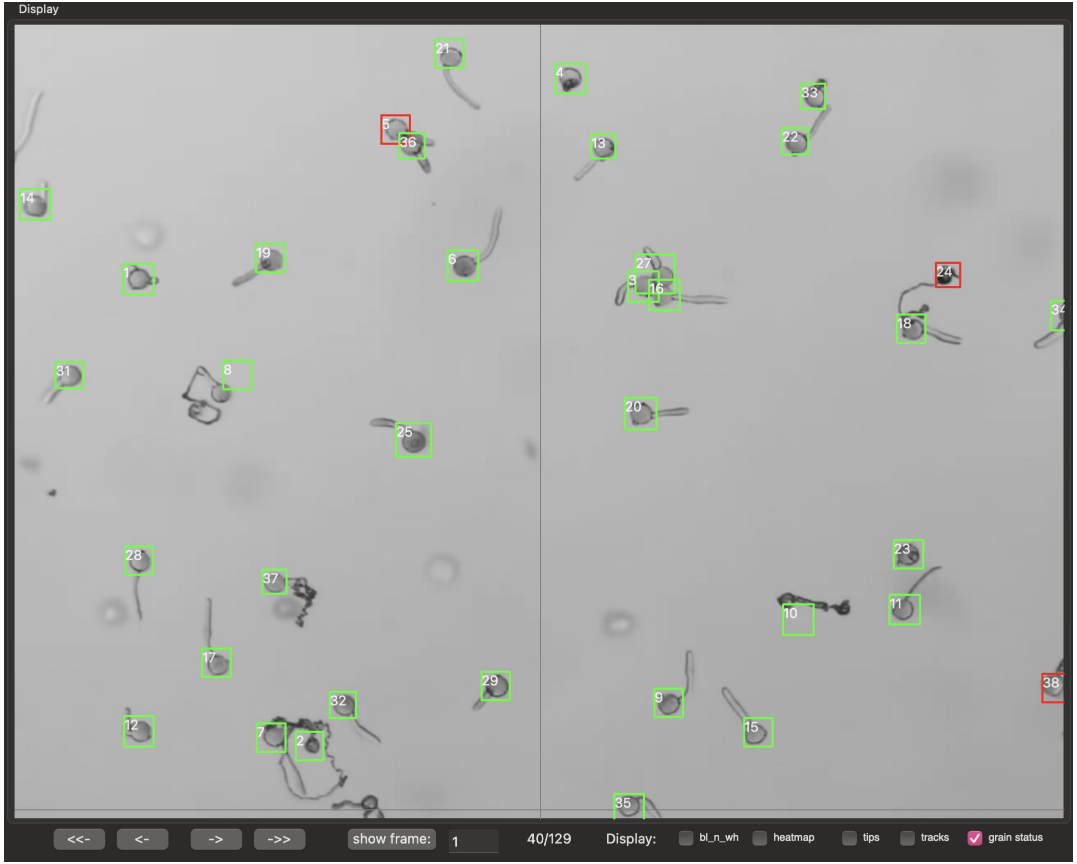


Using the **“Tip ovlp”** algorithm, taking 5 minutes to add undetected tips in early frames every 5 frames and applying the hint bellow, the majority of germination events are properly called (bounding box changes from red to green).

Hint: if tracking both grain germination and tip elongation, I would recommend tracking tip elongation first. At the end of this tracking, you will be asked if you would like to keep tip locations that were predicted to close gaps between tracks. More tip locations should improve the accuracy of the **“Tip ovlp”** algorithm.

### Manual Tracking and QC of germination

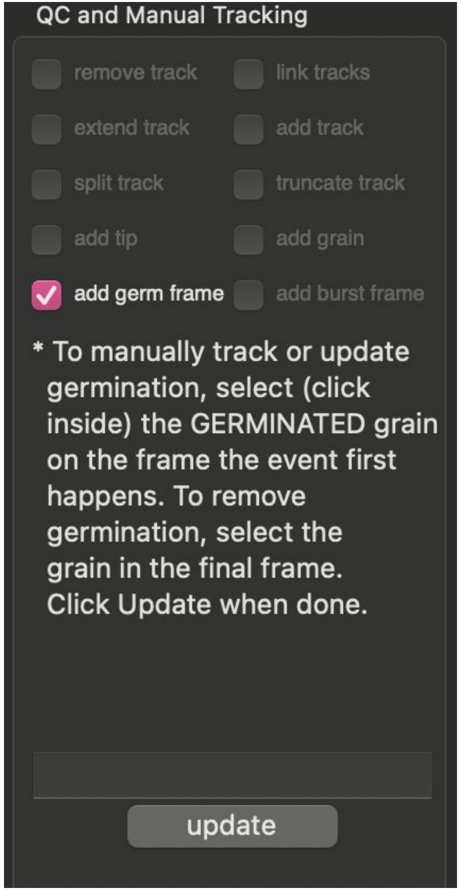


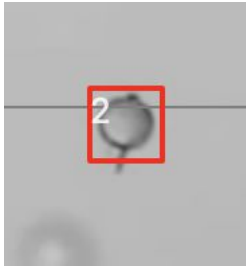


Grain 2 has germinated, but it was not detected

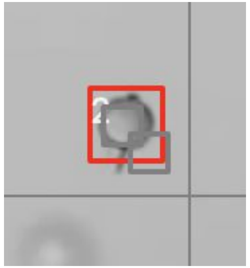

The user indicates the frame when germination occurred. if the same grain is selected multiple times, the most recent selection is considered

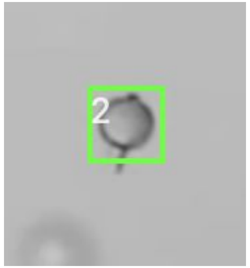


The bounding box changes to green for all subsequent frames

### Manual Tracking of pollen survival, an example

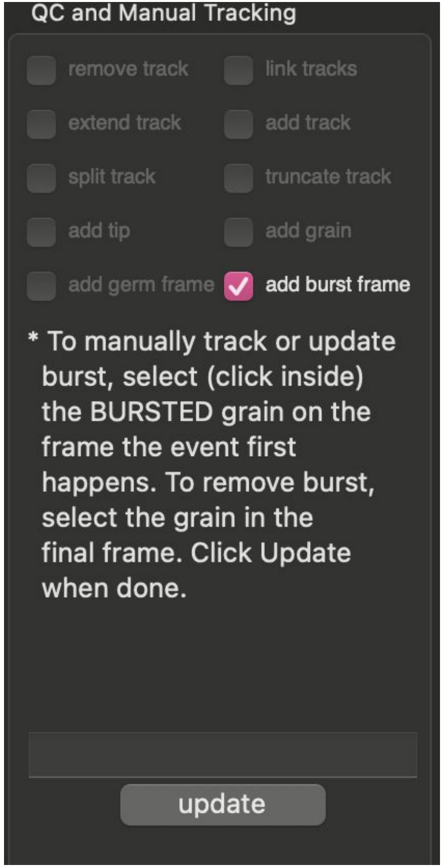


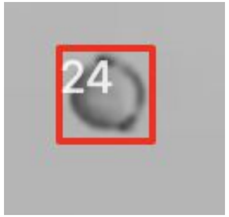


Grain 24 has not germinated (red bounding box)

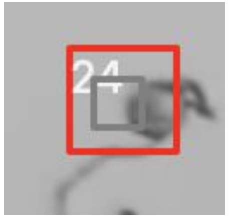

The user indicates the frame when grain burst occurs

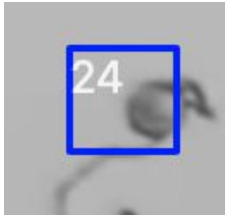


The bounding box changes to blue color for all subsequent frames

### Manual Tracking of pollen survival, an example

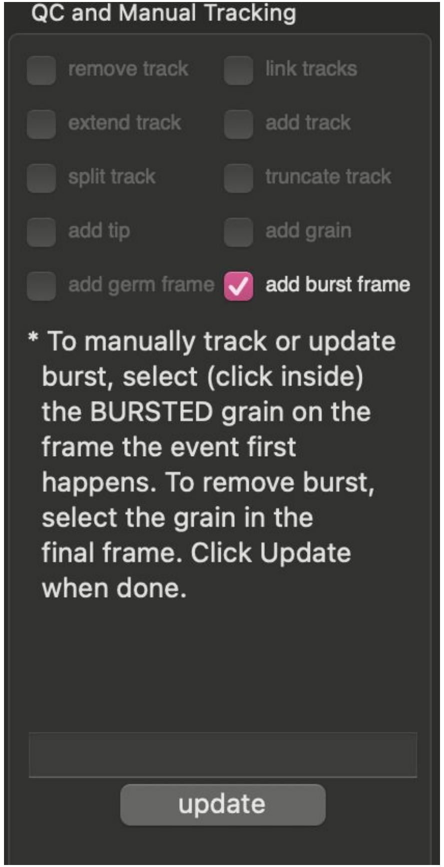


TubeTracker currently facilitates tracking pollen survival manually.

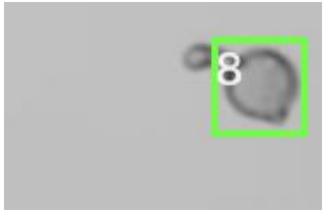


Grain 8 has germinated

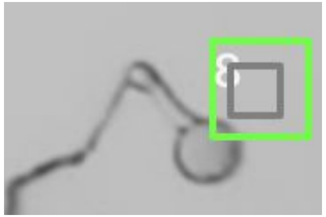


The user indicates the frame when pollen tube burst occurs

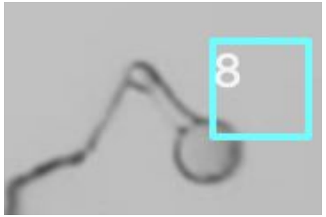


The bounding box changes to cyan color for all subsequent frames

### Usage Instructions

#### Saving results

### Saving results

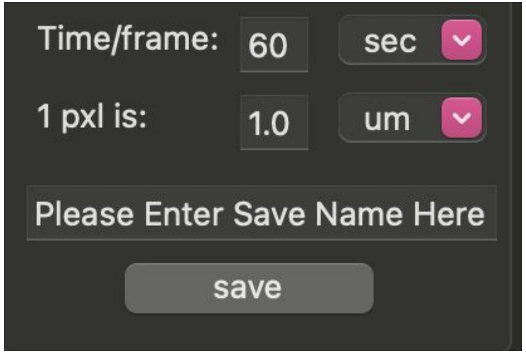


After tracking is complete, data can be exported into comma separated files (csv). Two videos summarizing tracking of 1) germination plus survival and 2) tip elongation are provided.

If the user tracked on germination/survival, but not tip elongation (or vice-versa), only the relevant files will be generated.

To generate results, the user must provide:

- **"Time/frame"**: The time between successive frames along with the time unit. Unit options include: sec, min, hour, day, week, month, year.
- **"1 pxl is"**: the actual distance corresponding to 1 pixel. Unit options include: um, nm, mm, cm, m, in, ft, pxl
- **"Saving name"**: the filename to use when saving files. If nothing is provided, "result" will be used as default.
- A directory to write the files into by clicking on **"save"**

### Saved files when tip elongation is tracked

- **result.tip.tracks.avi**: video summary of tip tracking.
- **result.tracks.raw.data.csv**: stores all raw data related to each track identified.

| track_id | frame | centroid_x | centroid_y | time (sec) | length (pixel) | length (um) | detection_method |
|---|---|---|---|---|---|---|---|
| 1 | 0 | 182 | 263 | 0 | 0 | 0 | auto |
| 1 | 1 | 181 | 262 | 60 | 1.8697580638061700 | 1.8697580638061700 | auto |

- **result.tracks.unsynchronized.csv**: reports lengths (user provided units) of converted tracks at each timepoint after detection

| time (sec) | track.1 length (um) | track.2 length (um) | track.3 length (um) | track.4 length (um) | track.5 length (um) | track.6 length (um) |
|---|---|---|---|---|---|---|
| 0 | 0 | 0 | 0 | | | |
| 60 | 2.907018061756960 | 1.285140562249000 | 1.285140562249000 | 0 | 0 | |
| 120 | 4.192158624005960 | 1.285140562249000 | 1.285140562249000 | 4.817263292661670 | 1.8697580638061700 | 0 |

- **result.tracks.synchronized.csv**: reports lengths (user provided units) of converted tracks with time 0 set to time of first detection.

| time (sec) | track.1 length (um) | track.2 length (um) | track.3 length (um) | track.4 length (um) | track.5 length (um) | track.6 length (um) |
|---|---|---|---|---|---|---|
| 0 | 0 | 0 | 0 | 0 | 0 | 0 |
| 60 | 2.907018061756960 | 1.285140562249000 | 1.285140562249000 | 4.817263292661670 | 1.8697580638061700 | 2.570281124497990 |
| 120 | 4.192158624005960 | 1.285140562249000 | 1.285140562249000 | 9.089413434757860 | 3.7395161276123400 | 5.140562248995980 |

### Saved files when grain germination and/or survival are/is tracked

- **result.survival.avi**: video summary of tracking.
- **result.survival.raw.data.csv**: stores all raw data related to each grain tracked.

| grain_id | detection_time (sec) | germination_time (sec) | burst_time (sec) | detection_frame | germination_frame | burst_frame |
|---|---|---|---|---|---|---|
| 1 | 0 | 240 | -1 | 0 | 4 | -1 |
| 2 | 0 | 780 | -1 | 0 | 13 | -1 |

| germination_p_value | centroid_x | centroid_y | area_at_detection | detection_method | germination_method | burst_method |
|---|---|---|---|---|---|---|
| 0.045000000000000000 | 118 | 244 | 784 | auto | tip_overlap | manual |
| 0.045000000000000000 | 280 | 680 | 900 | auto | tip_overlap | manual |

Can be used to quantify % germination and time of germination for each grain

Note: -1 is a default value to indicate the event was not recorded

- **result.survival.curves.csv**: stores the number and fraction of grains germinated and burst at each timepoint.

| frame | time (sec) | num_germinated | fraction_germinated | num_bursted | fraction_bursted |
|---|---|---|---|---|---|
| 0 | 0 | 0 | 0.0 | 0 | 0.0 |
| 1 | 60 | 0 | 0.0 | 0 | 0.0 |
| 2 | 120 | 0 | 0.0 | 0 | 0.0 |
| 3 | 180 | 1 | 0.03225806451612900 | 0 | 0.0 |

Provides a summary of % germination and burst over time.
